# The Influence of Shoreline Residential Development on Riparian and Littoral Habitats in the Belgrade Lakes of Maine

**DOI:** 10.1101/2024.08.06.606825

**Authors:** Catherine R. Bevier, F. Russell Cole, Manuel Gimond, Bruce F. Rueger

## Abstract

Residential development of lake shoreland can be accomplished with minimal negative impact to riparian and littoral habitat complexity, biological diversity, lake water quality, and general ecosystem health by using landscaping best practices. Preserving a heterogeneous and stratified vegetated buffer along the shoreline helps reduce erosion and nutrient runoff. In our study, we compared characteristics of undisturbed reference, developed but buffered, and developed but unbuffered properties along shorelines of three lakes in the Belgrade Lakes region of south central Maine, USA. Features of riparian habitats associated with reference and buffered developed sites were generally more similar to each other than to unbuffered developed sites. Significantly less shading along the shoreline, fewer trees and tall shrubs in buffer areas, and narrower buffer strips along the shoreline occurred on unbuffered developed sites compared to buffered developed sites. The degree of substrate embeddedness, aufwuchs cover, and amount of woody structure were more similar for littoral habitats adjacent to undeveloped reference and buffered developed sites than for unbuffered developed sites. These findings represent important validation for the use of landscaping best management practices that mimic natural landscape patterns to minimize the influence of shoreland residential development on lake ecosystem health. These practices should be promoted by lake protection programs and communicated to shoreline property owners.

---

Lakes provide essential ecosystem services on global, regional, and local scales, and contribute billions of dollars of revenue as freshwater resources and recreational centers (Boyle et al. 1997, Bouchard 2000, Schuertz et al. 2001). However, development near freshwater ecosystems is negatively affecting ecosystem health (e.g., Carpenter et al. 1998) and the National Lakes Assessment reported that only 53% of the 111,818 lakes included in the study are in “a least disturbed condition” (US EPA 2016). Disturbed or degraded shoreland is the primary cause for lakes not achieving this standard, and for related declines in water quality and biological diversity in the adjacent lake ecosystems (Boyle et al. 1999, Jennings et al. 1999, Jennings et al. 2003, Brauns et al. 2007b, Brauns et al. 2011).

The state of Maine has 5,785 lakes covering roughly 11,766 km^2^ (Hasbrook 1995), and these lakes generate over $3.5 billion in economic value, support 52,000 jobs, and provide approximately 400,000 Mainers with clean drinking water (Boyle et al. 1997, Boyle et al. 1999, Bouchard 2000, Schuertz et al. 2001, NRCM 2013). More than 60% of Maine residents participate in lake-related recreational opportunities annually (MDEP 2017a). Protecting these valuable natural resources and mitigating the ecological impacts of changing shoreland use, especially residential development, have become important priorities for government agencies, lake managers, and lake community stakeholders.

Maine recognized early the importance of these natural resources by establishing some of the most progressive shoreland zoning laws in the country, including The Mandatory Shoreland Zoning Act of 1971 (38 MRSA §435–449) and the Natural Resource Protection Act of 1987 (38 MRSA §480). These establish protected zones and minimum setback requirements to promote lake ecosystem health (MDEP 2008, Merrell et al. 2013). These laws have been effective in preventing the degradation of aquatic shallow-water habitats and their biological diversity, and in providing sufficient protections to support the ecological integrity of existing aquatic communities (Merrell et al. 2013).

Growth of state-wide conservation programs such as the Lake Stewards of Maine (formerly Maine Volunteer Lake Monitoring Program, MVLMP 2017) and LakeSmart (Welch and Smith 2008, MLS 2016, Cole et al. 2017, Shannon et al. 2017), the formation of hundreds of citizen-based lake associations and conservation organizations, and the funding for extensive academic and government research on lakes have also helped to protect the health of Maine lakes. Even with these efforts, twenty-one of Maine’s lakes, including Great Pond in the Belgrade Lakes region, are listed as impaired by the US EPA (MDEP 2022), and other lakes are experiencing a rapid decline in water quality. In essence, Maine residents and visitors may be “loving their lakes to death” through unsustainable practices.

There is ample evidence of negative impacts occurring in riparian and shallow-water littoral habitats attributed to proximal residential development compared to habitats with natural, undeveloped landscape (e.g., Ness 2006, Carpenter et al. 2011, Merrell et al. 2013). Few studies, however, document the link between specific onshore riparian habitat that reflect important buffering capacities, and features of the near shore littoral environment. A well-maintained buffer area is an important component of effective lake protection programs, and should be integrated into all lake management strategies (Merrell et al. 2009, 2013).

In this study, we document how unbuffered and buffered shoreland residential development influence abiotic and biotic characteristics of riparian and littoral environments for three lakes in south central Maine. Given that disturbance or degradation of shoreland can negatively affect lake ecosystem health (e.g., Carpenter et al. 1998, Kaufmann et al. 2014), we hypothesized that abiotic and biotic characteristics of riparian and littoral environments adjacent to developed, but effectively buffered properties, across the three lakes studied would not differ significantly from those of undeveloped reference lots compared to unbuffered properties, and that the latter would be more prone to erosion and runoff of sediment and nutrients, and less suitable for wildlife than undeveloped or well-buffered, developed properties. Support for these hypotheses provides validation for lake protection programs that promote the use of landscaping best management practices and confirms that the presence and maintenance of intact and well-vegetated, multilayered buffers enable residential development to occur on shoreland with minimal degradation to lake ecosystem health.

## METHODS

### Study area

The Belgrade Lakes, located in south central Maine, are an interconnected chain of seven major lakes (Fig. 1) and an important driver of the local Maine economy (Boyle et al. 1997, Bouchard 2000, Schuertz et al. 2001). They were formed as the last continental ice sheets passed southward through Maine and scoured out their basins, which are underlain by igneous and metamorphic rock (Hildreth 2005, Thompson 2015). Glacial retreat deposited sands and gravels in the basins, and these sediments were then redistributed by wave activity to create the current environment (Table 1).

**Figure 1.**
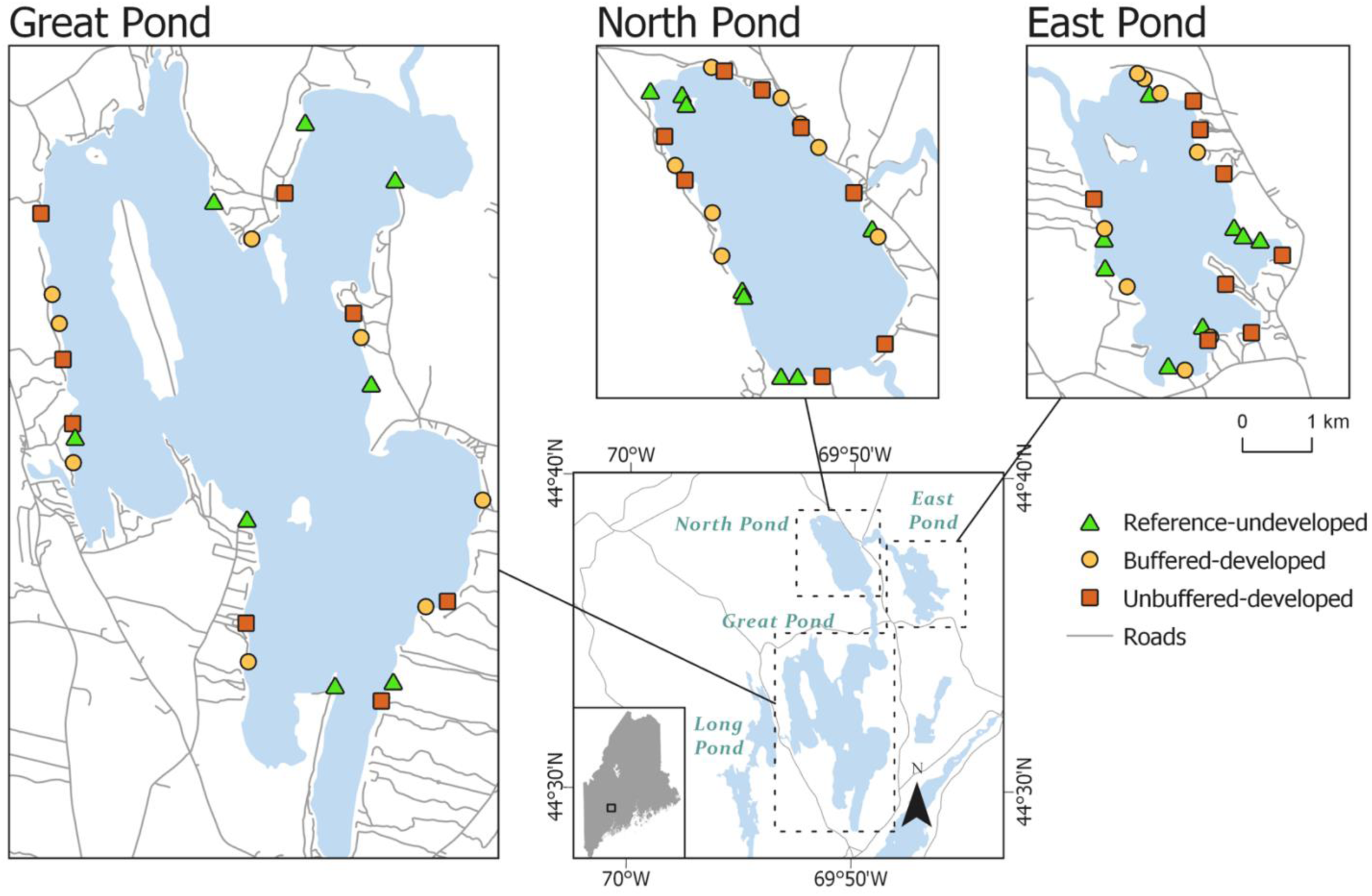
Map of the Belgrade Lakes region located in south central Maine, USA showing the relative location of the three study lakes: East Pond, North Pond, and Great Pond. The distribution of shoreline sampling sites REF (Reference, undeveloped), BD (Buffered, developed), and UD (Unbuffered, developed) is shown for each of the study lakes. The scale bar applies to the three lake maps and not the inset map.

**Table 1.**
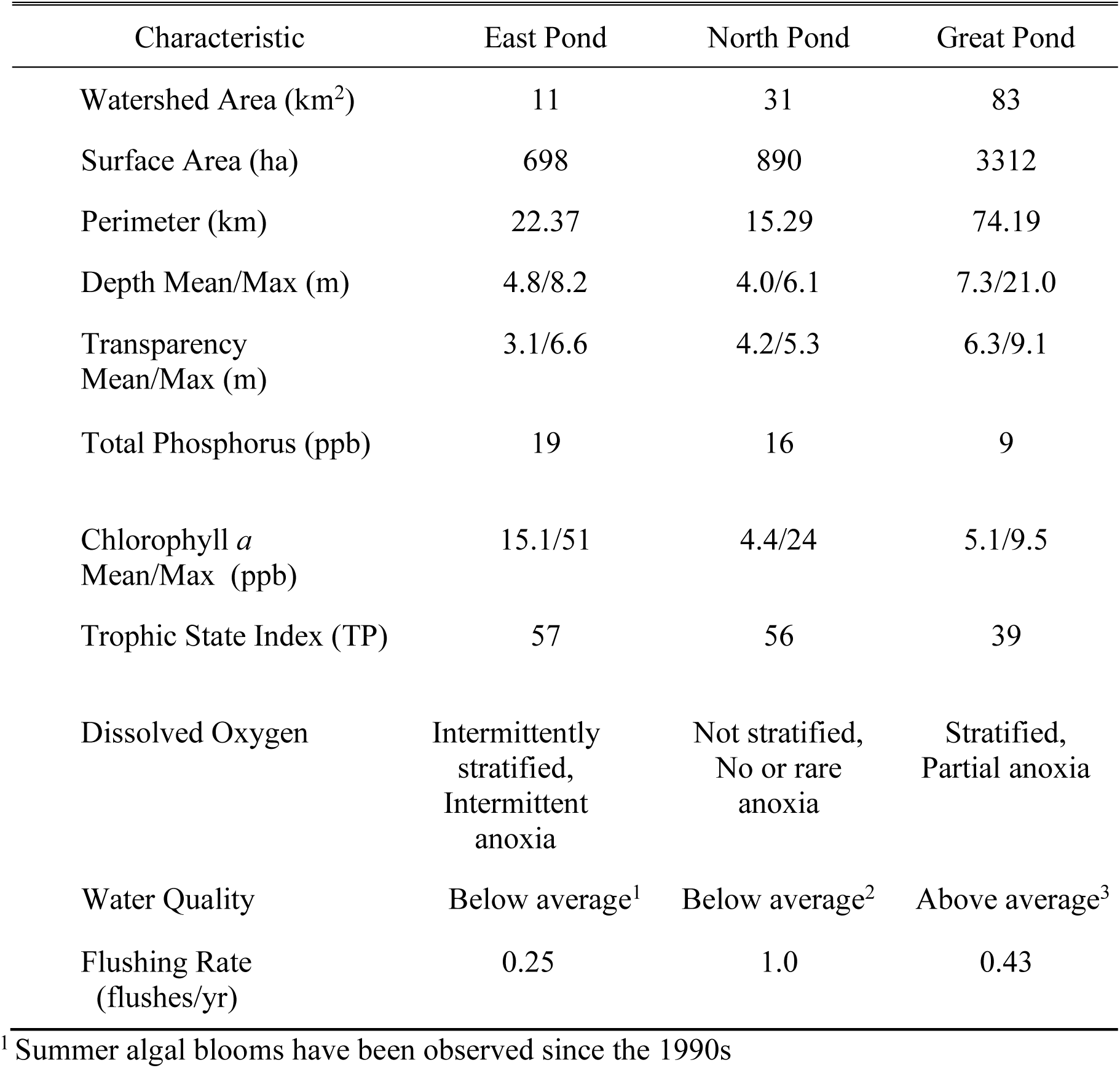
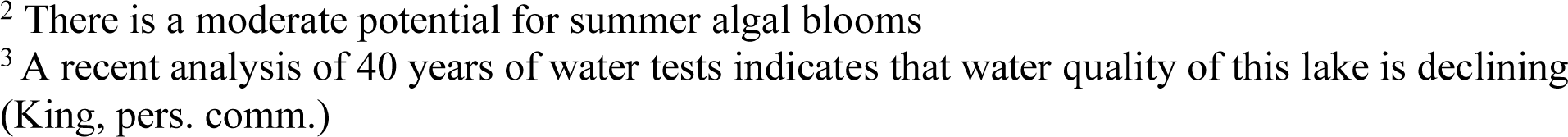
Characteristics of East Pond, North Pond, and Great Pond located in the Belgrade Lakes area of south central Maine, USA. We provide physical and water quality assessments for these lakes obtained from the Maine Department of Environmental Protection (MDEP 2017b) and the Volunteer Lakes Monitoring Program (MVLMP 2017). The trophic state index for each lake was based on total phosphorus concentrations and calculated following (Carlson 1977).

| Characteristic | East Pond | North Pond | Great Pond |
| --- | --- | --- | --- |
| Watershed Area (km<sup>2</sup>) | 11 | 31 | 83 |
| Surface Area (ha) | 698 | 890 | 3312 |
| Perimeter (km) | 22.37 | 15.29 | 74.19 |
| Depth Mean/Max (m) | 4.8/8.2 | 4.0/6.1 | 7.3/21.0 |
| Transparency<br>Mean/Max (m) | 3.1/6.6 | 4.2/5.3 | 6.3/9.1 |
| Total Phosphorus (ppb) | 19 | 16 | 9 |
| Chlorophyll *a*<br>Mean/Max (ppb) | 15.1/51 | 4.4/24 | 5.1/9.5 |
| Trophic State Index (TP) | 57 | 56 | 39 |
| Dissolved Oxygen | Intermittently stratified, Intermittent anoxia | Not stratified, No or rare anoxia | Stratified, Partial anoxia |
| Water Quality | Below average<sup>1</sup> | Below average<sup>2</sup> | Above average<sup>3</sup> |
| Flushing Rate (flushes/yr) | 0.25 | 1.0 | 0.43 |

<sup>1</sup> Summer algal blooms have been observed since the 1990s

Over the last five decades, the Belgrade Lakes region has changed considerably from a landscape historically dominated by agricultural land use, low population density, clear healthy lakes, and seasonal residences scattered along the lake shoreland to a patchwork of fields and forests inhabited by a higher human population density and dominated by mesotrophic lakes. Our study focused on shoreline properties located on East Pond (towns of Oakland and Smithfield) in Somerset County, North Pond (also known as North Lake and Little Pond; towns of Mercer, Rome, and Smithfield), and Great Pond (towns of Belgrade and Rome) both in Kennebec County.

While these ponds share a similar geological history which influenced their physical formation and general characteristics (Thompson 2015), they differ in size, shape, and bathymetry as well as the degree of shoreland residential development, the mean age of the shoreline residences, and the ratio of seasonal to year-round residences along the shoreline (Table 1, unpublished data). The catchments of these lakes differ in size and are dominated by rolling hills covered by fallow fields and mixed hardwood and evergreen forests (the six most common tree species in rank order are *Tsuga canadensis*, *Acer rubrum*, *Pinus strobus*, *Betula papyrifera*, *Fraxinus* sp., and *Quercus rubra*) interspersed with scattered patches of fertile farmland. These lakes are surrounded by older seasonal cottages interspersed with an increasing number of newer year-round residences. Many of the older residences (pre–1971) were constructed on or near the waterline; post–1971 residences must adhere to the setback regulations and are located at least 100 ft from the water’s edge. Small rural communities are scattered throughout the entire Belgrade Lakes catchment.

The Lake Stewards of Maine (formerly Maine Volunteer Lake Monitoring Program, MVLMP) provided Secchi disk measurements, chlorophyll *a,* and total phosphorus concentrations for each of the three study lakes (Table 1, MVLMP 2017). The trophic state index for each lake was based on total phosphorus concentrations obtained from epilimnetic core samples taken at the deepest point in the lake and calculated following Carlson (1977).

### Study design

Our methodology parallels that of Kaufmann and Whittier (1997) and Merrell et al. (2009, 2013), who developed protocols for rapid evaluation of abiotic and biotic habitat characteristics (e.g., substrate size and type, littoral habitat for biota, macroinvertebrate abundance and diversity, riparian vegetation cover and structure) for near-shore riparian habitat and shallow-water littoral habitat. We used a balanced sampling design for this study that included equal representation of three residential development “treatments” on each of three study lakes in the Belgrade Lakes catchment (Fig. 1). These lakes share similar environmental and exposure conditions. All sites were visually identified from a lakeside view and selected to maximize consistency within each treatment type.

The three treatment categories used in this study were undeveloped (reference properties = REF), developed properties with a well-formed vegetated buffer (buffered developed = BD), and developed properties without a buffer (unbuffered developed = UD). Reference sites had intact habitat consisting of stratified layers of ground cover, low and tall shrubs, trees along the property shoreline, no evidence of recent disturbance or erosion, and no built structures. Buffered developed properties included at least one built structure, a heterogeneous and stratified vegetated buffer along the shoreline, and exhibited little evidence of erosion or the presence of erosion prone areas. Unbuffered developed properties were extensively landscaped, often with large impervious lawn areas running to the shoreline with little to no vegetated buffer along the shoreline.

UD and BD treatments included both seasonal and year-round residences, some built close to the shoreline (pre–1971) and others set back from the shoreline following current MSZA regulations (MSZA, 38 MRSA §435–449; NRPA, 38 MRSA §480). With permission from the owners, we surveyed eight properties for each of the three treatment types on each of the three study lakes. We used a clustered design for each replicate where properties representing each of the three treatment types were roughly proximate to one another. However, this arrangement was not always possible given the actual distribution of undeveloped, buffered developed, and unbuffered developed properties along the shoreline. On each lake, treatment clusters were located to reflect all directional points of exposure (Fig. 1).

### Survey: Riparian Habitat

At each treatment site, we estimated the degree of shoreline disturbance (i.e., what proportion of the shoreline appeared natural versus altered or removed). We scored the observed disturbance as low (1 = < 32%), medium (2 = 33-66%), or high (3 = > 67%). We evaluated shoreline stability (i.e., what proportion of the shoreline appears effectively buffered against wave-caused erosion) as low (1 = < 32%), medium (2 = 33-66%), or high (3 = > 67%). We measured the slope (%) of the property from a canoe located in front of the property and close to the shore using a clinometer. We also recorded the presence of large lawn areas and noted areas of significant erosion.

To quantify riparian vegetation type and cover, we used a 10 m x 10 m quadrat centered between property boundaries with the lower edge placed on the shoreline (Fig. 2). Within the quadrat, we determined the mean, maximum, and minimum buffer width for each property by measuring the narrowest and widest points in the buffer (up to the 10 m quadrat boundary). We also estimated the relative cover of each riparian vegetative component (i.e., groundcover, low shrubs, tall shrubs, and trees) within the quadrat. Because these components were evaluated independently, the percent cover represents individual density estimates for each vegetation component found in the quadrat and may exceed 100%. The ground cover component included long grass, broad leaf weeds, duff, moss, leaf litter, or rock. We noted when maintained lawn was present within the quadrat, but did not consider it as part of natural ground cover. We estimated riparian vegetation complexity using a composite score (0–12, representing low to high), calculated by summing the cover score (0 to 3 to reflect 0%, ≤ 50%, and ≤ 100%) for each of the four vegetation components (i.e., groundcover, low shrubs, tall shrubs, and trees). We identified each tree in the quadrat to species, measured its diameter at breast height (DBH), classified it as regenerating (< 10 cm DBH), young (11–28 cm DBH), or mature (> 29 cm DBH), and counted the total number of trees per species. Finally, we measured percent shading by vegetation along the riparian shoreline standing on land at 1.0 m from the water’s edge and centered on the 10 m nearshore quadrat boundary, and standing in the water in the same centered position on the 0.5 m and 1.0 m depth contours using a densiometer. Percent shading was calculated for each of the three sampling locations following Lemon (1956).

**Figure 2.**
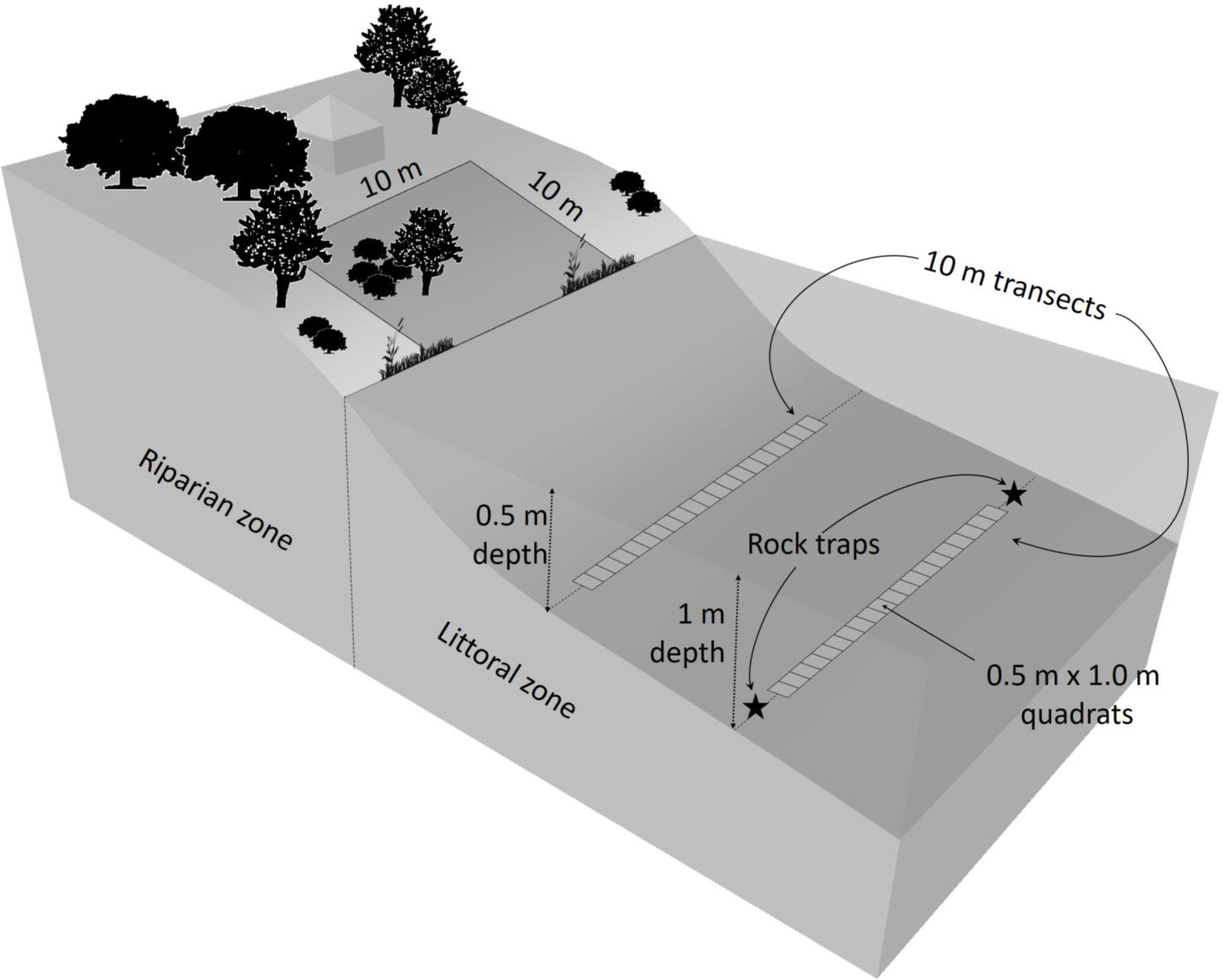
Diagram of the sampling design at an individual study site. The riparian habitat was sampled using a 10 m x 10 m quadrat. The littoral habitat was sampled using 0.5 m and 1.0 m depth transects with twenty 1.0 m x 0.5 m quadrats located along each transect. The asterisks at the ends of the 1.0 m depth transect indicate the location of rock traps.

### Survey: Littoral Habitat

We characterized the littoral environment at each treatment site by first placing a 10 m weighted transect line marked at 1.0 m increments along the 0.5 m depth and 1.0 m depth contours (Fig. 2). Although the nearshore transect (0.5 m depth) was most directly influenced by the adjacent terrestrial ecosystem, we included the 1.0 m depth transect to help ensure that we investigated a broad range of the possible shallow-water littoral benthos. We used 0.5 m x 1.0 m quadrats made of PVC pipe to quantify littoral substrate characteristics. Each quadrat was subdivided by sturdy cord to form 50 squares (each 2 % of the quadrat area). Twenty of these quadrats were centered perpendicularly and placed side by side on the submerged transect line to create a survey band along the entire 10 m length of each transect. Surveyors moved slowly along each transect to survey each of the 20 quadrats. Surveyors used viewing buckets with clear plexiglass bottoms to eliminate glare off the water surface to improve visibility through the water column to the substrate to facilitate the survey. Surveyors estimated the percent cover for different substrate types (i.e., boulders, cobble, gravel, sand, and silt) along the length of each transect. We estimated the degree of embeddedness using protocols from Platts et al. (1983) and Ness (2006), which were based on percent cover of large benthic rocks by fine sediments or sand; the embeddedness categories used were < 5%, 5–25%, 25–50%, 50–75%, and 75–100%.

In addition to these visual transect and quadrat surveys, we conducted sediment core sampling during summer 2013 using a custom core sampler to collect samples from the top 5 cm of substrate from three randomly selected locations along each of the 0.5 m and 1.0 m depth transects at each treatment site. We placed the collected samples in Whirl-pak® bags and kept these samples on ice until we returned to the laboratory where they could be transferred to a freezer until analyzed. To determine the distribution of sediment grain sizes, we dried samples in ceramic crucibles for 24 hours at 65° C, then poured them into a 2 mm sieve. We removed and weighed the large pebbles (> 2 mm) to comprise one sediment sample component, then weighed remaining particles that were less than 2 mm in size to comprise the second sample component. The total sediment weight for a sample equaled the sum of these two components. The size distribution of particles < 2 mm was determined by placing roughly 100 g of sediment into an automatic shaker holding a set of 10 sieves of different mesh sizes (1.40 mm, 1.00 mm, 0.710 mm, 0.500 mm, 0.355 mm, 0.250 mm, 0.180 mm, 0.125 mm, 0.090 mm, and 0.063 mm) and shaking the sieves for 20 min. The weight of the contents of each of the 10 sieve mesh sizes was recorded to determine the relative distribution of sediment sizes. Finally, to conduct an analysis of ash-free dry mass (AFDM), we placed approximately 3 g of < 2 mm particle sediment from each core sample in a crucible and heated it to 105° C for 12 hours to remove all moisture. We then burned the samples in a muffle furnace at 550° C for 12 hours and recorded the difference between initial and final weights to obtain AFDM, and we report the mean percent organic matter for each site.

We visually determined a cover score for aufwuchs, the community of algae, bacteria, fungi, protozoans, micro- and macroinvertebrates attached to the substrate surfaces in the littoral environment, by estimating the proportion of rock surface within a quadrat that was covered. For each of the twenty 0.5 m x 1.0 m quadrats placed along the 10 m transect, we determined an aufwuchs cover score using a scale of 1–10, with 1 representing 10% of total rock surface covered and 10 representing 100% coverage, and calculated a mean for the twenty quadrats. We estimated aufwuchs density using a scale of 1 to 3 (sparsely to densely covered) for each quadrat, then calculated a mean for the twenty quadrats that comprised the 10 m x 1.0 m band transect.

We quantified combined deciduous and coniferous leaf litter cover present on littoral substrate by counting the number of quadrat squares covered for each of the twenty 0.5 m x 1.0 m quadrats (Fig. 2). These values were summed over the length of the 10 m band transect and converted to percent cover by multiplying the sum by 0.1 to estimate the percent of litter present along the entire transect. We also counted the pieces of coarse (>10 cm diameter), medium (4–10 cm), and fine (< 4 cm diameter) woody structure in each quadrat and recorded a total for each category for the 20 quadrats located along the 10 m x 1.0 m band transect.

We sampled macroinvertebrate diversity along the 0.5 m and 1.0 m depth transects using two different methods. First, surveyors identified single locations at the midpoint of each transect, then shoveled the top layer of substrate, including rocks, sticks, and detritus, from roughly 0.25 m^2^ into a 500 µm Surber net, then emptied and rinsed the net into a bin of water. We scrubbed any rocks present into the bin then returned them to the pond. We used a 600 µm sieve to strain the collected organisms out of the bin water. We preserved the invertebrates collected in bottles containing 70% ethanol. We also sampled macroinvertebrate diversity and recruitment into the littoral habitat of treatment sites using rock traps. Each rock trap (32 cm long x 9 cm diameter, volume of 8143 cm^3^) was filled with clean rocks, ranging from golf ball to tennis ball size. One rock trap was placed at each end of the 1.0 m depth transects (Fig. 2). We collected the rock traps after three weeks, and followed protocol described above for cleaning rocks, and collecting and preserving captured specimens.

Invertebrates collected using both sampling procedures were identified, typically to genus or family, using a combination of dichotomous keys and online resources (e.g., Peckarsky 1990, Merritt et al. 2008). We calculated a Shannon Index for collections from each treatment site to compare overall macroinvertebrate diversity across treatments on each lake and among the three study lakes. We investigated the diversity of taxa considered tolerant of organic pollution by determining the family-specific feeding habits as defined by Mandaville (2002). The Family-level Biotic Index (FBI) was then calculated using the products of the number of organisms in each family and their familial tolerance values, summing these products, and dividing the sum by the total number of organisms collected (Hilsenhoff 1988).

In addition to estimating pollution tolerance, we used the percentage of the total number of organisms classified as Crustacea and Mollusca (freshwater clams), taxa known to be sensitive to pollution, as a separate metric, and used the same approach for individuals from the Orders Coleoptera, Odonata, Trichoptera, and Ephemeroptera (COTE) as an indicator of good water quality. These metrics served as indicators of good water quality and overall lake health (Brauns et al. 2007b, De Sousa et al. 2008) and were chosen to eventually allow a comparison of results to those reported for a similar study in Vermont (Deeds, pers. comm.). We also used the proportion of Chironomidae and Oligochaeta as indicators of disturbance (Brauns et al. 2007b, De Sousa et al. 2008).

### Statistical analysis

We completed statistical analyses using R (R Core Team 2018). We assessed variation in the measured variables for the riparian and littoral habitats of the three study lakes (East Pond, North Pond, Great Pond) and used development treatments of properties (REF, BD, UD) as the nominal variables. We used non-parametric statistical tests because the data were not normally distributed (Shapiro Wilk test). We applied a Kruskal-Wallis test using p < 0.05 to determine if the mean ranks of a measured variable were different among lakes, regardless of treatment within a lake, and among treatments across lakes. P-values were adjusted for multiple comparisons using Holm’s adjustment to correct for family-wise error rate (Holm 1979). If at least one of the treatment groups was significantly different for a particular variable, we followed up with pair-wise comparisons of the variable across lakes or treatment types with a Dunn test (Dunn 1964) as implemented in the R package *FSA* (Ogle 2018).

We also explored trends across treatment groups using the Jonckheere-Terpstra rank-based trend test (Terpstra 1952, Jonckheere 1954) whereby the treatments were ranked in the order of reference, buffered and unbuffered. This test looks for an ordered pattern to the median of the groups being compared and was implemented using R’s *clinfun* package (Seshan 2018). Because of ties in the data, the package’s built-in permutation option was used to compute an estimate of the exact *P* value. Percent organic matter in sediment samples was analyzed by first transforming data, found to be highly heteroskedastic, using the inverse of the values. We then assessed the effect of season and lake interactions, then applied a one-way ANOVA. Supplementary tables with mean values (± 1 SE) for all variables and with outcomes of statistical tests are available online.

## RESULTS

### Riparian Habitat

#### Across Lakes

The percent of residences surveyed that met the 100 ft set back regulation differed across lakes: East Pond had only one property with a structure set less than 100 ft from the water. Great Pond and North Pond properties had a higher proportion of this type of structure location; 25% and 29%, respectively, had structures that did not adhere to this regulation. Properties on the three lakes were similar in nearly all variables measured to evaluate riparian habitat, including shading over onshore and nearshore habitat from overhanging vegetation, and the widths of the riparian buffer measured within our quadrats (Fig. 3, Appendix S1: Table S1). Approximately two thirds of the properties assessed on Great Pond (67%) and East Pond (63%) possessed riparian buffers that exceeded the 10 m width of our quadrat compared to 21% of the North Pond riparian buffers.

**Figure 3.**
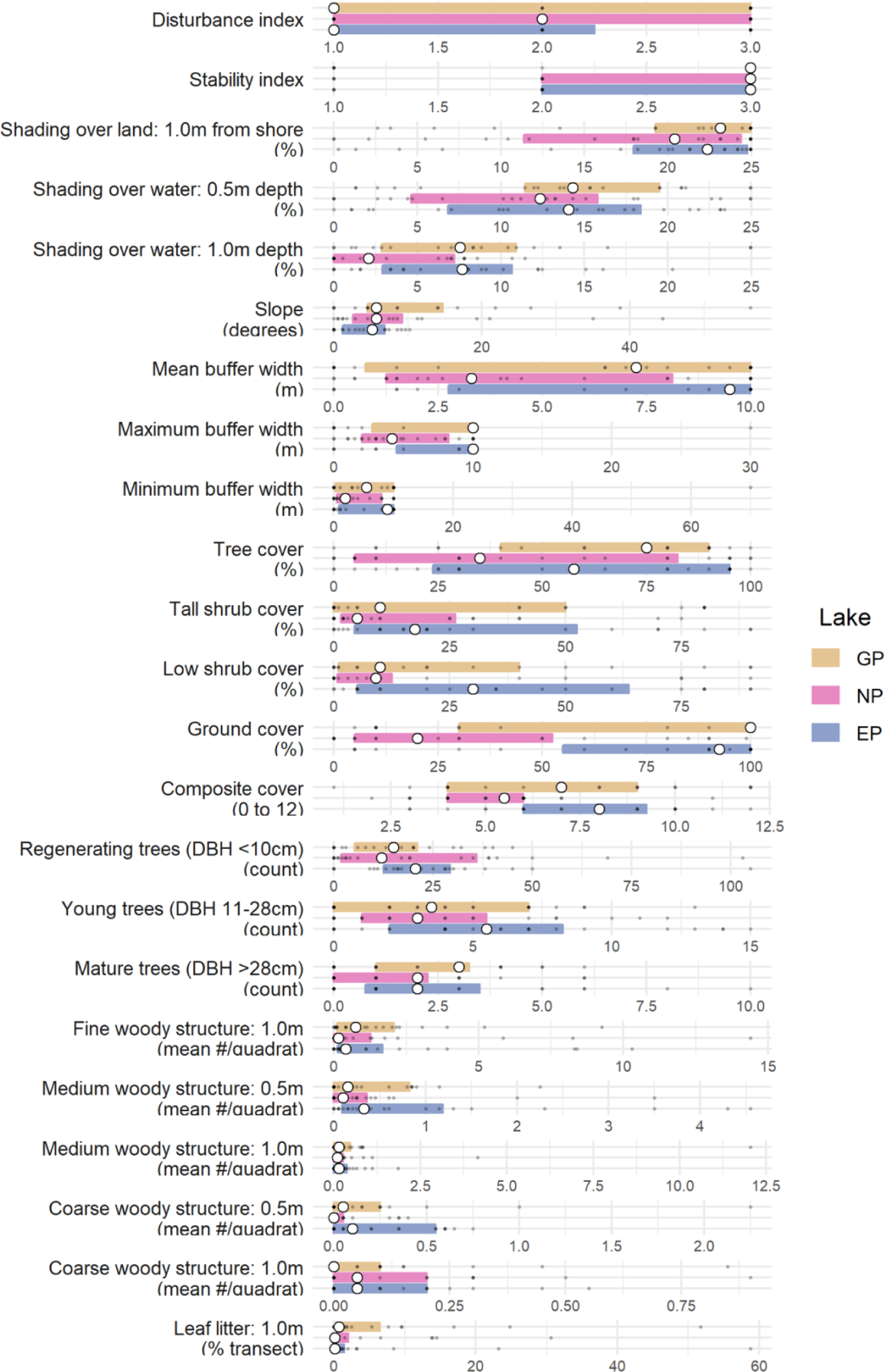
Summary plots of riparian measurements aggregated by lake (GP = Great Pond, NP = North Pond, EP = East Pond). Colored boxes encompass the interquartile ranges. The open circles represent the sample medians and the smaller gray filled circles represent individual values.

The percent vegetation cover in the 10 m x 10 m riparian quadrats composed of trees and shrubs did not differ among properties on different lakes, but the percent groundcover vegetation identified was significantly different (Kruskal-Wallis; H = 17.25, *P* = 0.003; Fig. 3, Appendix S1: Table S1); properties on North Pond had statistically lower percentages of groundcover compared to East and Great Ponds (Appendix S1: Table S1). Large expanses of lawn area that could facilitate sediment, nutrient, and contaminant runoff were present on 22% of all sites surveyed, with some variation across lakes (33% of the North Pond sites, 21% of the East Pond sites, and 10% of the Great Pond sites). Finally, the number of trees within the quadrats for each of our three tree size categories was not statistically different across the three lakes (Fig. 3, Appendix S1: Table S1).

#### Across Treatments

In contrast to our comparisons across lakes, most of the variables in the riparian habitat differed significantly across treatments. The shoreline disturbance by human activity, as measured by a disturbance index, was statistically different across treatments within a lake (Kruskal-Wallis test; H = 54.21, *P* < 0.001; Fig. 4, Appendix S1: Table S2). BD and REF shorelines were similar, but both treatments were roughly one-half and one-third as disturbed as UD shorelines. The index of shoreline stability also differed across treatments (Kruskal-Wallis test; H = 22.74, *P* < 0.001); REF and BD properties were similar, but shorelines of UD properties were statistically less stable than either REF or BD properties (Fig. 4, Appendix S1: Table S2). Mean slope of the riparian area was greatest for REF sites and least for UD sites, but differences across the three treatments were not statistically different (Fig. 4, Appendix S1: Table S2).

**Figure 4.**
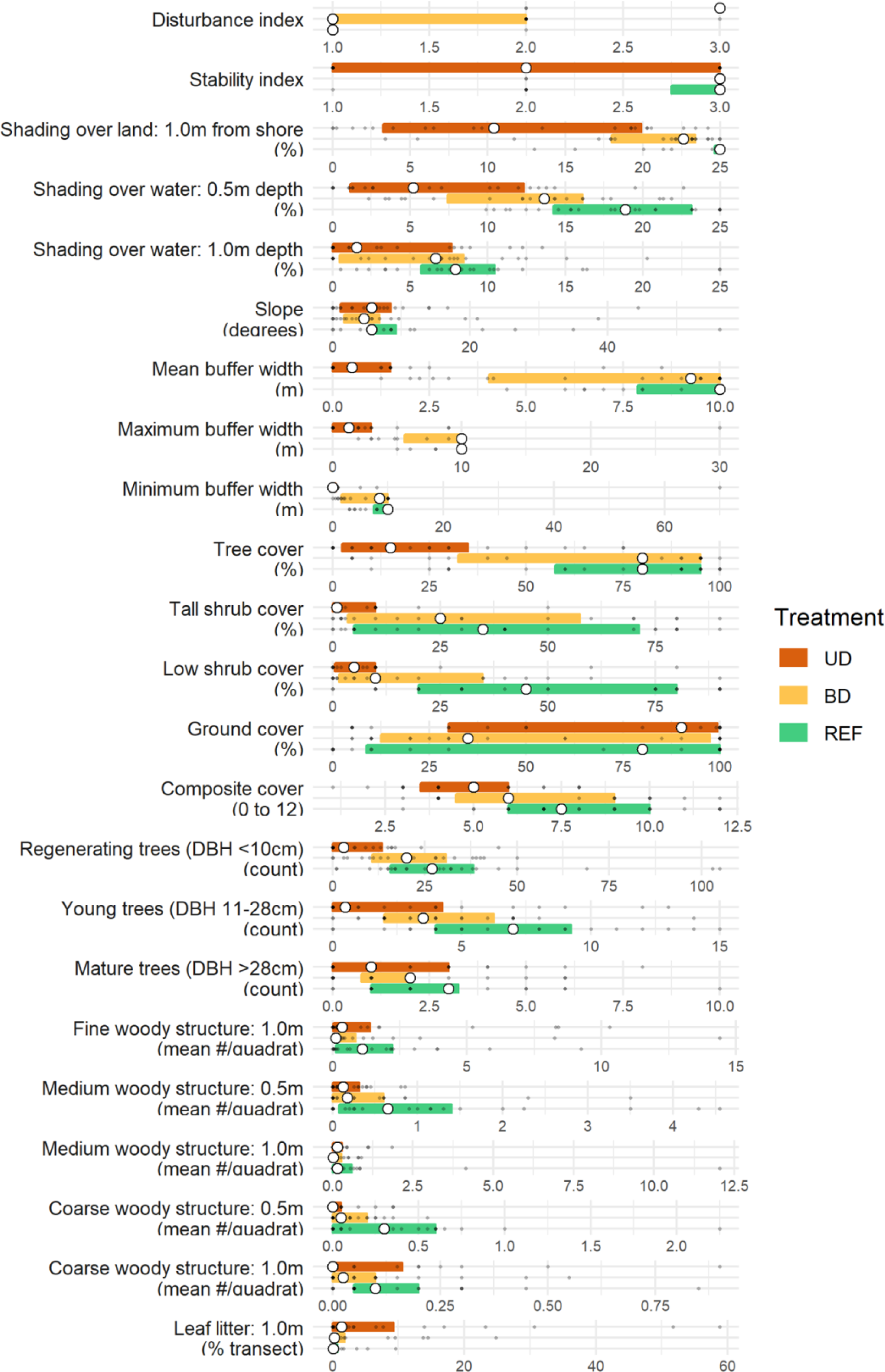
Summary plots of riparian measurements aggregated by treatment type (UD = Unbuffered, developed, BD = Buffered, developed, REF = Reference). Colored boxes encompass the interquartile ranges. The open circles represent the sample medians and the smaller gray filled circles represent individual values.

The percent vegetation shading on land and at both transect depths was lowest for UD sites and overall more similar between REF and BD sites, but differed statistically when compared across all treatments (Kruskal-Wallis test; H = 34.0, 25.64, 10.30, respectively for measured locations at REF, BD, and UD sites; *P* < 0.001, <0.001, 0.023). REF properties had the highest percent vegetation shading and UD sites had the lowest shading at all measurement locations (Fig. 4). BD sites were statistically more shaded than UD sites over land and shallow water, and REF sites were statistically more shaded than BD sites over land and shallow water (Fig. 4, Appendix S1: Table S2). REF and UD sites were statistically different for shading measurements over land and at both water depths (Fig. 4, Appendix S1: Table S2).

The differences in mean, maximum, and minimum widths of riparian buffer areas measured within our quadrats were statistically different across the three treatment types (Kruskal-Wallis test; H = 43.40, 39.33, 35.54, respectively for REF, BD, and UD sites; *P* < 0.001 for all; Fig. 4), with all three measures being statistically lower for UD properties than for REF and BD properties (Appendix S1: Table S2). Mean and maximum riparian buffer widths were not statistically different between BD and REF sites, but REF sites had statistically greater minimum width than BD sites (Fig. 4, Appendix S1: Table S2). The riparian buffer extended beyond the quadrat boundary (10 m) for the majority of REF (83%) and BD (61%) properties. In comparison, no UD property possessed a riparian buffer that extended beyond the 10 m x 10 m sample quadrat.

Statistical differences in percent vegetation cover estimates for trees, tall shrubs, and low shrubs recorded from the quadrats, and the vegetation cover composite index calculated from these component estimates, were observed across treatments (Kruskal-Wallis test; H = 24.24, 18.81, 18.63, 17.08, respectively; *P* ≤ 0.001 for all). These metrics were highest for REF and BD sites, but significantly lower for UD sites compared to BD and REF (Fig. 4, Appendix S1: Table S2). The composite index and percent cover for trees and tall shrubs did not differ statistically between REF and BD sites, and the percent vegetation cover for the low shrub component was statistically higher for REF sites than for either BD or UD sites (Appendix S1: Table S2). Although the estimate of ground cover as a percentage of the 10 m x 10 m composition was somewhat higher for UD sites than for REF and BD sites, likely because of the more open canopy at UD sites, these estimates were not statistically different. Large areas of relatively impervious lawn were most commonly found on UD sites (52% of those surveyed) whereas none of the REF properties and 12.5% of the BD properties included lawn among components of groundcover. More pervious areas with duff and long grass were dominant components of groundcover, and taller vegetation such as trees and shrubs composed a much greater proportion for REF and BD properties compared to UD properties.

The number of regenerating and young trees differed across the treatments (Kruskal-Wallis test; H = 22.0, 13.93, respectively; *P* < 0.001, 0.005; Fig. 4, Appendix S1: Table S2). REF and BD sites had similar numbers of small regenerating trees, but both treatment types had at least three times more small regenerating trees than the UD sites (Appendix S1: Table S2). REF sites had statistically more young trees than either BD or UD site, and BD sites had statistically more young trees than UD sites (Appendix S1: Table S2). The number of mature trees was low and similar at all treatment sites (Fig. 4, Appendix S1: Table S2).

### Littoral Habitat

#### Across Lakes

Overall, the near shore littoral habitats were similar across the three lakes studied, differing only in the composition of the benthic substrate measured as percent cover of each substrate type along a 10 m x 1.0 m transect band at both 0.5 m and 1.0 m depths (Fig. 5). Statistical differences in the percent sediment cover of boulders and gravel were observed along the 0.5 m depth transect (Kruskal-Wallis test; H = 29.10, 19.38; p < 0.001, *P* = 0.001 respectively), and the percent cover of boulders observed along the 1.0 m depth transect (Kruskal-Wallis test; H = 24.65; *P* < 0.001). Statistically more boulders were found along the transects at both 0.5 m and 1.0 m depths in East Pond than along transects in either North Pond (Wilcoxon rank-sum tests; *P* < 0.001 for both depths) or Great Pond (Wilcoxon rank-sum tests; *P* < 0.001, *P* = 0.002, respectively), which did not exhibit statistically significant differences (Fig. 5). Great Pond had statistically higher percent gravel cover along the 0.5 m depth transect than either East Pond or North Pond, which had considerably lower gravel cover (Wilcoxon rank-sum tests; *P* = 0.01, < 0.001, respectively; Fig. 5). While the difference among lakes was not significant at the corrected significance value, East Pond and North Pond had higher percent silt cover on the substrate than Great Pond (Fig. 5).

**Figure 5.**
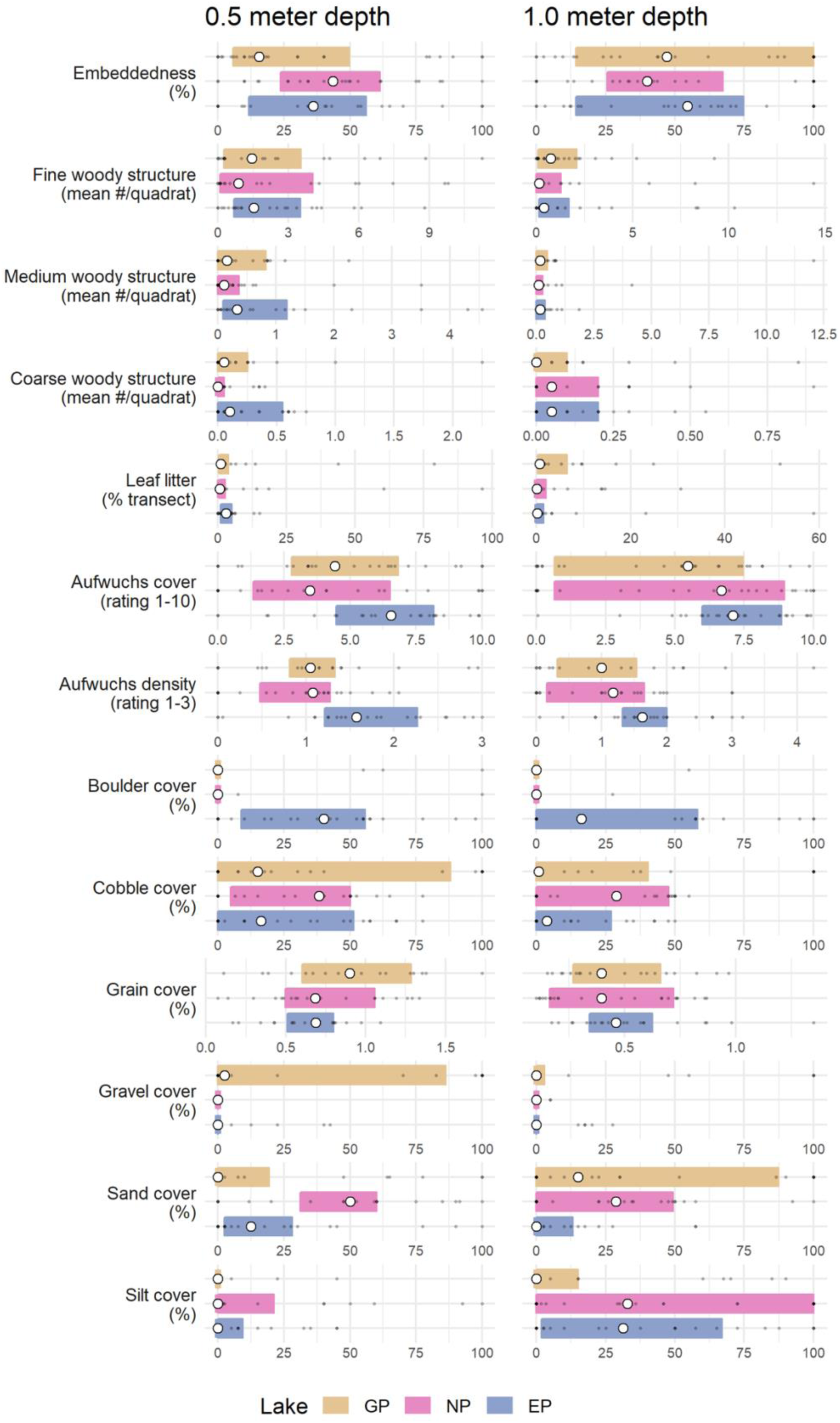
Summary plots of littoral measurements aggregated by lake (GP = Great Pond, NP = North Pond, EP = East Pond). Colored boxes encompass the interquartile ranges. The open circles represent the sample medians and the smaller gray filled circles represent individual values.

Differences in the organic component of the sediments, reported as a percent of the total dried and ashed sample mass, collected and analyzed early in the summer were statistically higher for East Pond than North Pond or Great Pond, which were similar (one-way ANOVA (F_(2, 64)_ = 10.622, *P* = 0.001; Appendix S1: Table S3). The late season samples did not differ across the three lakes, and there was no interactive effect of time of sampling. Percent embeddedness of benthic rocks was similar across the three lakes along both transect depths (Fig. 5, Appendix S1: Table S3). The accumulated fine, medium, and coarse woody structure observed along the 0.5 m and 1.0 m depth transects were generally similar across the three lakes studied, as were the aufwuchs cover and density, and leaf litter cover (Fig. 5).

We assessed macroinvertebrate diversity across the three study lakes using data from rock traps and from grab samples. There were statistical differences in total number of organisms, mean taxon richness, Family-level Biotic Index (FBI), and proportion of Chironomidae among lakes based on data from rock traps (Kruskal-Wallis test; H = 16.97, 13.34, 25.44, 31.33; *P* = 0.003, 0.019, < 0.001, and < 0.001 respectively; Fig. 6a, Appendix S1: Table S4). Sites in Great Pond yielded significantly fewer total organisms and lower mean richness of taxa than either East Pond or North Pond (Appendix S1: Table S4). FBI and percentage of invertebrates represented in the Family Chironomidae, the latter being an indicator of disturbance, were highest for East Pond and lowest for North Pond (Fig. 6a, Appendix S1: Table S4). Overall, mean taxon richness was higher and more total organisms were collected by grab sampling in all three lakes along the 1.0 m depth transects than along the 0.5 m depth transects, although these differences were not statistically different for any of the lakes, and there were no differences in any of the metrics assessed across lakes at either depth (Fig. 6b, Appendix S1: Table S4).

**Figures 6a, b.**
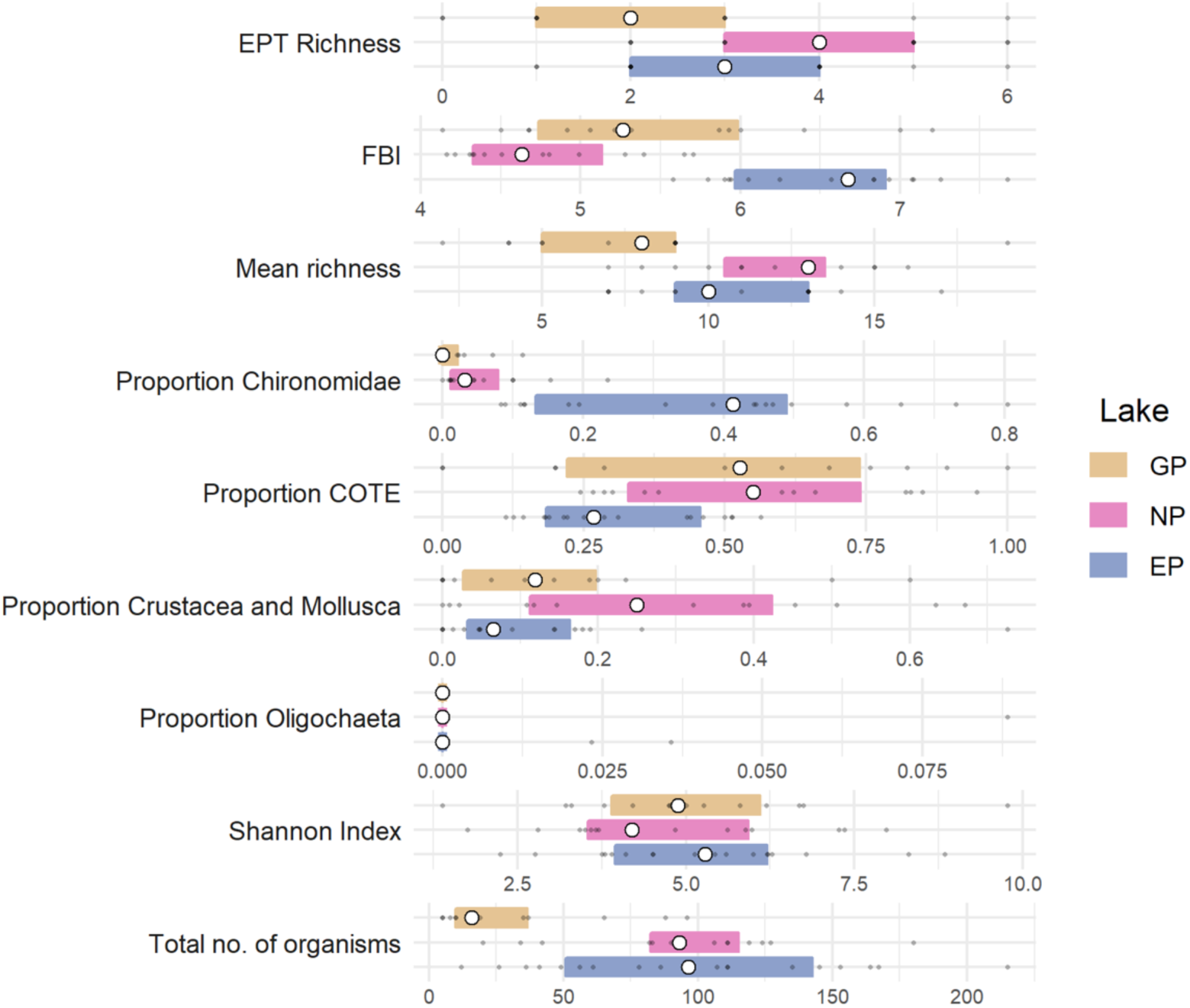
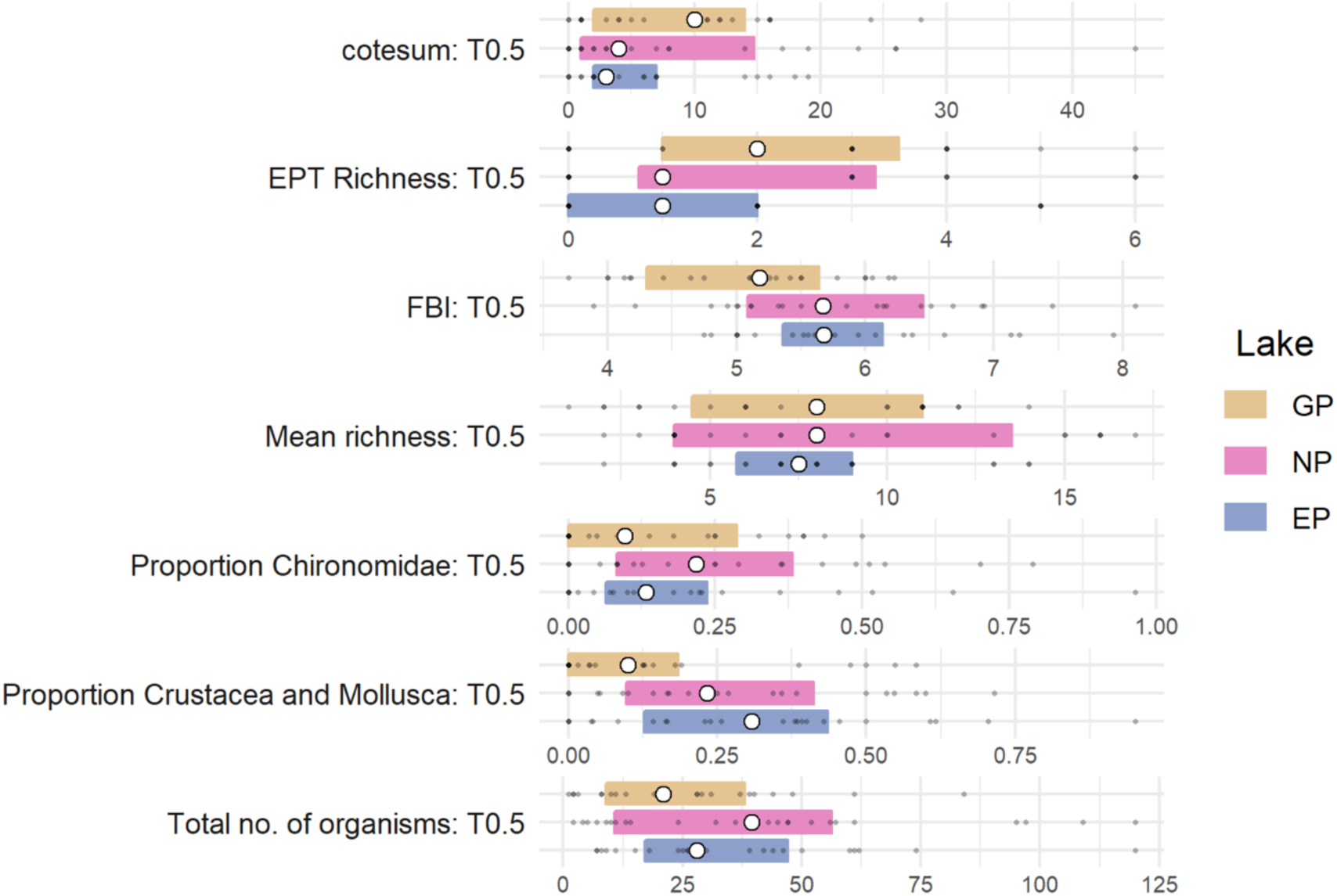
Summary plots of invertebrate diversity evaluated using rock traps (a) and grab samples along a transect, aggregated by lake (GP = Great Pond, NP = North Pond, EP = East Pond). Colored boxes encompass the interquartile ranges. The open circles represent the sample medians and the smaller gray filled circles represent individual values.

#### Across Treatments

The differences in percent cover of boulders, cobble, gravel, and silt in the littoral substrate at 0.5 m and 1.0 m depths were not statistically different across the three treatments (Fig. 7). However, the differences in percent sand in the substrate along the 0.5 m depth transect were statistically significant (Kruskal-Wallis test; H = 16.9, *P* = 0.006). UD sites had an almost three times higher percentage of the substrate composed of sand (52%) than for REF (20%) and BD (17%) sites (Wilcoxon rank-sum tests, *P* < 0.001 for both). REF and BD sites had similar and low levels of sand along the 0.5 m transect (Fig. 7), and all three treatments were similar for this metric along the 1.0 m water depth transect. Embeddedness of large rocks in the substrate did not differ across treatments (Fig. 7, Appendix S1: Table S5). The organic composition of the sediments was higher for REF sites than UD or BD sites, but this difference was not statistically significant for either sample season (Appendix S1: Table S5).

**Figure 7.**
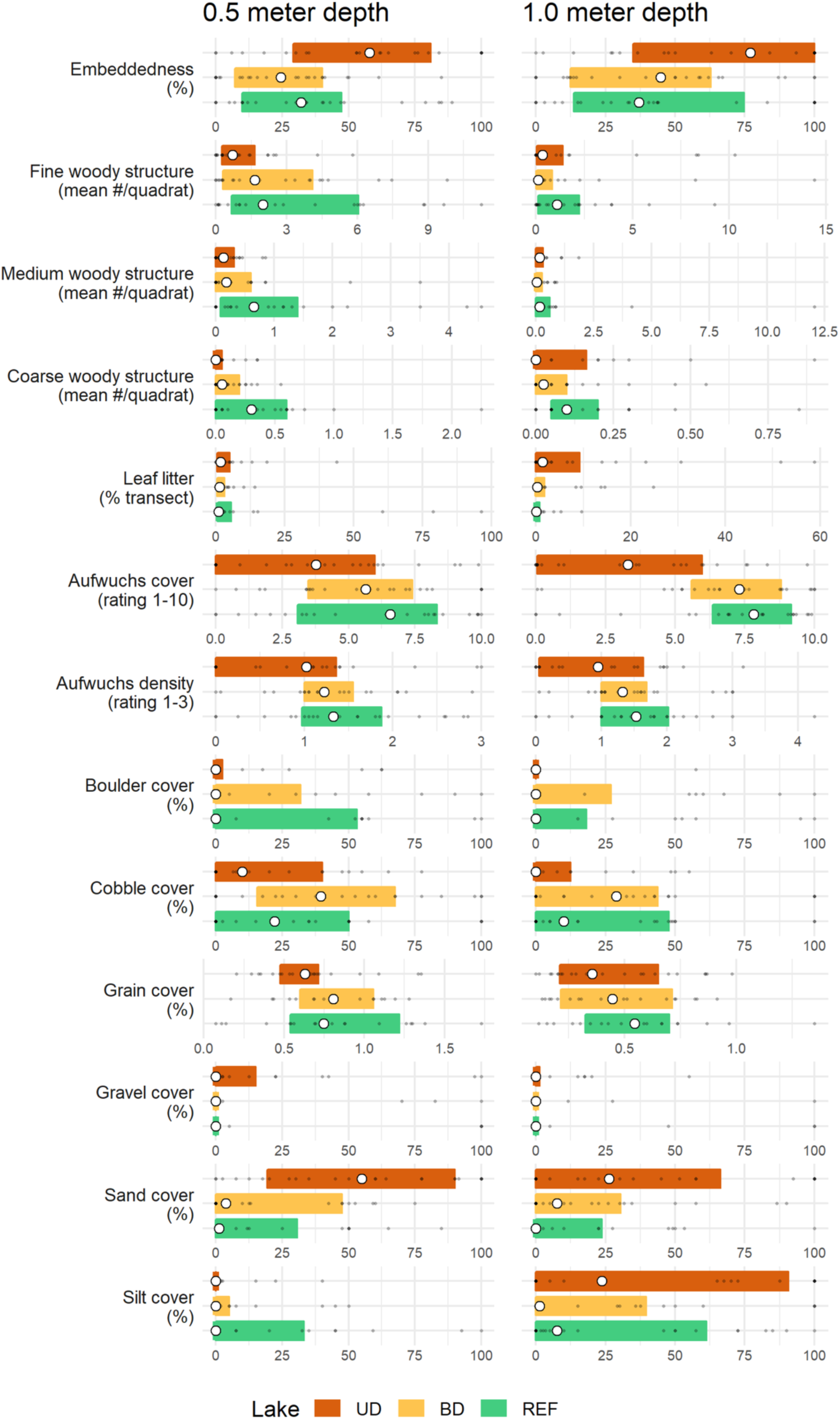
Summary plots of littoral measurements aggregated by treatment type (UD = Unbuffered, developed, BD = Buffered, developed, REF = Reference). Colored boxes encompass the interquartile ranges. The open circles represent the sample medians and the smaller gray filled circles represent individual values.

The number of pieces of woody structure in all three size-classes and leaf litter on the benthic surface were similar across the three treatment types at both transect depths (Fig. 7). Although the REF sites had more pieces of fine, medium, and coarse woody structure than the BD and UD sites along both transects, the only statistical difference was in coarse woody structure along the shallow water transect (Fig. 7, Appendix S1: Table S6). Mean rating for aufwuchs cover scored at 1.0 m depth transects was statistically different across treatments (Kruskal-Wallis test; H = 16.63, *P* = 0.006) and lower for UD sites than BD and REF (Fig. 7, Appendix S1: Table S6). No statistical differences were detected for aufwuchs density or macrophyte abundance at either 0.5 m and 1.0 m transect depth across the three treatment types.

Rock trap samples evaluated across treatments were similar, but total number of organisms collected using grab samples differed significantly across treatment sites along the 0.5 m depth transects (Kruskal-Wallis test; H = 14.81, *P* = 0.013; Fig. 8a, Appendix S1: Table S7); BD sites had nearly twice as many organisms as REF and UD sites. None of the metrics were statistically different across the three treatments for samples collected along the 1.0 m depth transects (Fig. 8b, Appendix S1: Table S7).

**Figures 8a, b.**
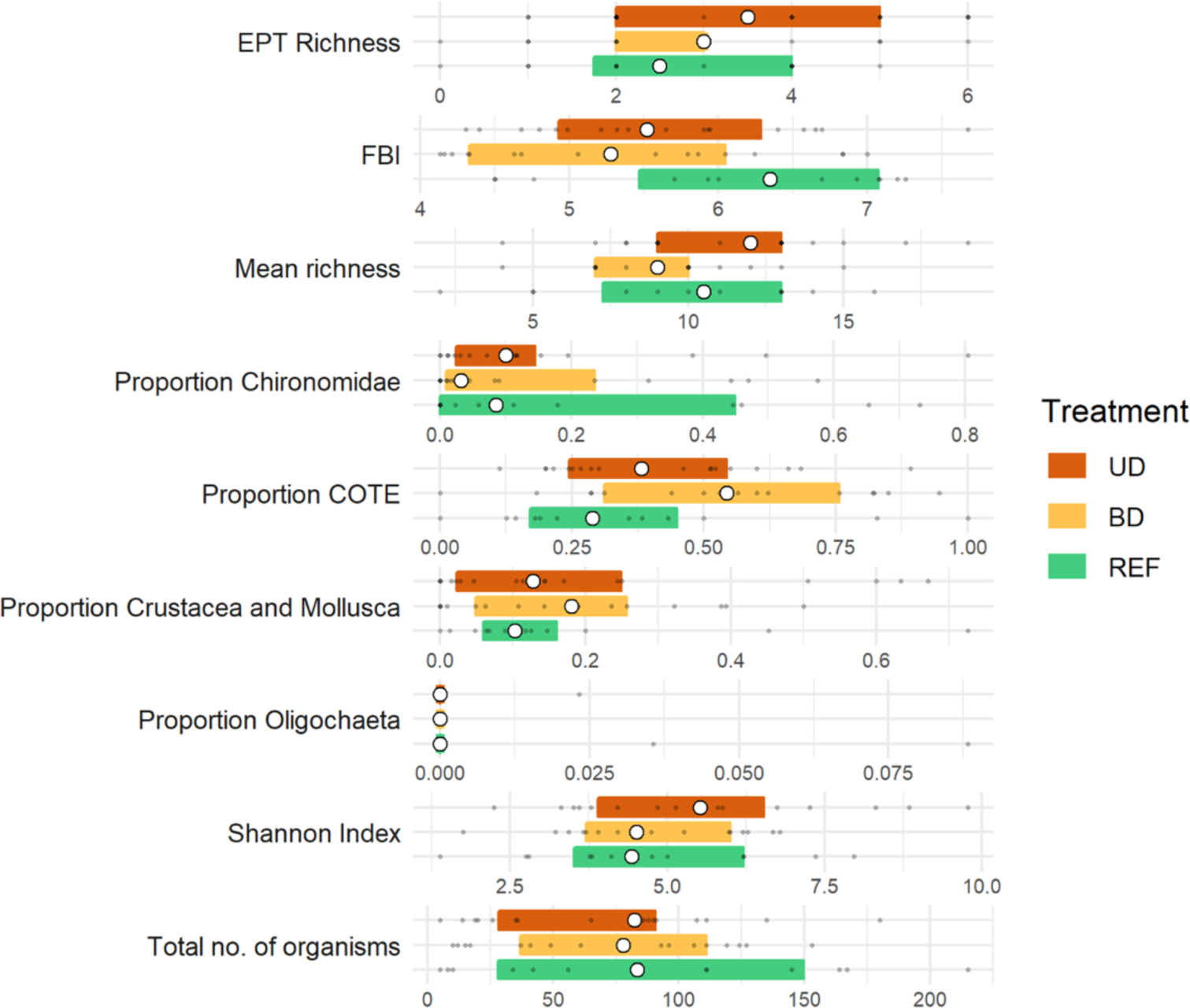
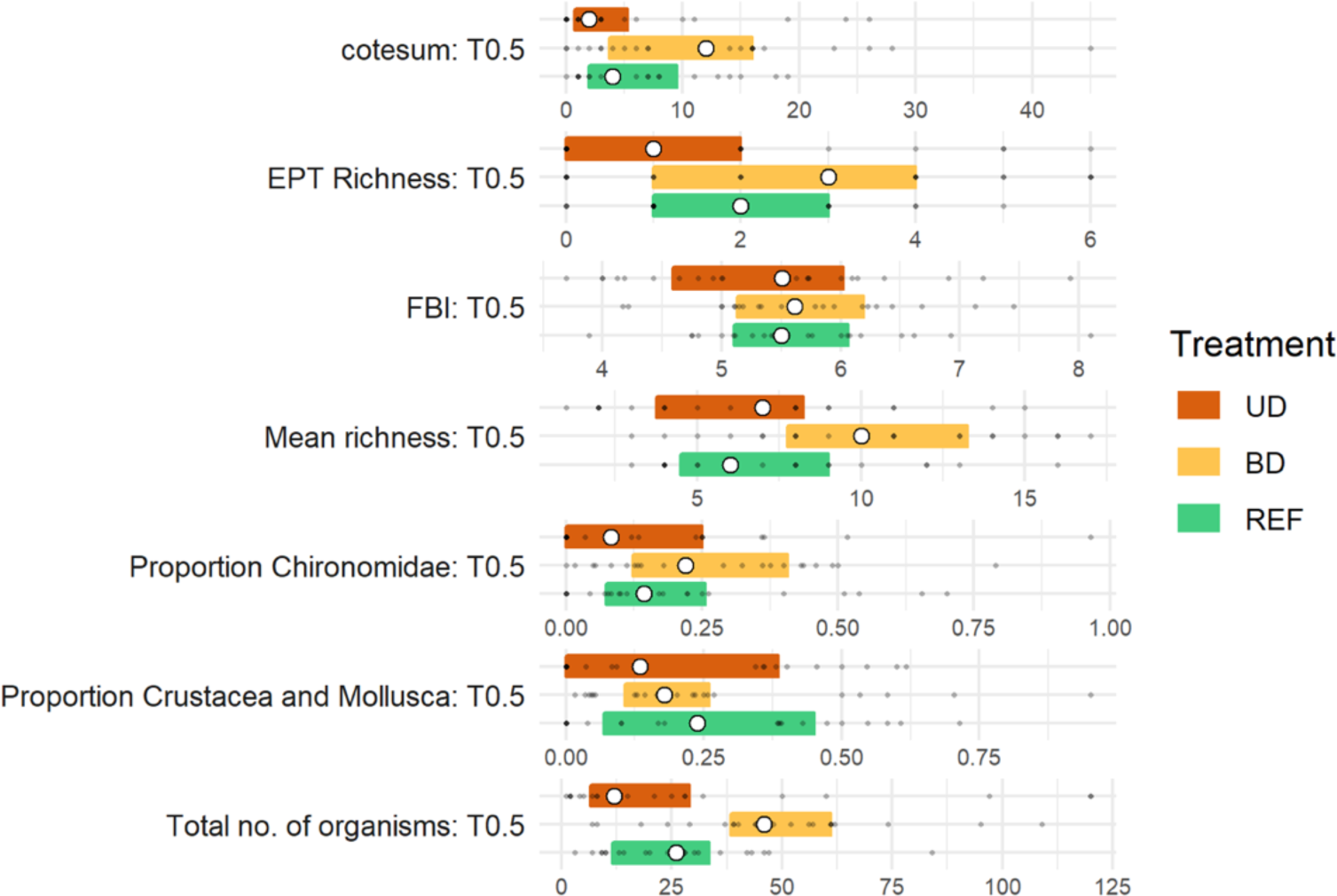
Summary plots of invertebrate diversity, evaluated using rock traps (a) and grab samples along a transect, aggregated by treatment type (UD = Unbuffered, developed, BD = Buffered, developed, REF = Reference). Colored boxes encompass the interquartile ranges. The open circles represent the sample medians and the smaller gray filled circles represent individual values.

## DISCUSSION

Unbuffered developed residential properties (UD) on the shorelands of East Pond, North Pond, and Great Pond in the Belgrade Lakes catchment in south central Maine lack a robust riparian buffer and cause disturbance and degradation to the adjacent shallow-water littoral habitat. Such properties had landscapes without trees and shrubs but with manicured, large, and relatively impervious lawns extending to the water’s edge. This kind of property management facilitates runoff to flow into the lake, especially during heavy rains, making the shoreline more vulnerable to erosion and nutrient runoff, less suitable for wildlife, and more likely to have a negative impact on lake ecosystem health than reference (REF) or buffered, developed properties (BD). Abiotic and biotic characteristics of riparian and littoral environments on or adjacent to buffered, developed (BD) properties, on the other hand, were similar and, in many cases, not statistically different from those of the riparian and littoral environments on or adjacent to REF sites. Most of the BD shoreline properties in this study had environmentally-friendly landscaping, including undisturbed and complex riparian vegetation composed of a mixture of grasses, low and tall shrubs, and both deciduous and coniferous trees. These effective riparian buffers promote filtration, deposition, and sorption of sediment, nutrients, and other contaminants before runoff reaches the lake, and help maintain the ecological integrity of the adjacent littoral environment and lake health in general (Barling and Moore 1994, Carpenter et al. 1998, Stutter et al. 2012, Allaire et al. 2015). The types of vegetation, the degree of complex stratification, buffer width and slope, and the location of trees and shrubs within the riparian habitat can influence the effectiveness of runoff absorption into the soil before the surface flow enters the lake (Welsch 1991, Hickey and Doran 2004, Liu et al. 2008, Zhang et al. 2010).

All three study lakes, however, have residences built before current zoning regulations were approved in 1971, and are therefore exempt from current legal restrictions that include a 100 ft set back. Without a riparian buffer, these properties can greatly influence lake ecosystem health, particularly for the two smaller and shallower lakes (East Pond and North Pond). The presence of old houses constructed near the shoreline and the resulting disturbance to and limitations for riparian vegetation may help explain the observed lower shading over the water (at 1.0 m depth), the lower percent of ground cover, and the lower composite vegetation cover index in the riparian areas of North Pond. The findings from this study validate the importance of constructing effective riparian buffers on developed properties when natural buffers are not present or have been largely degraded. Maintaining these buffers over time, especially for those properties with structures located close to the shoreline is also important. We found that BD sites were more similar to the REF sites than UD sites in many characteristics of riparian areas generally. For example, riparian buffers at the BD sites provided more shading on land and over water, and were more than four times wider and more similar to the REF sites than the buffers present on the more disturbed UD sites. Percent vegetation cover of trees and tall shrubs, which helps to mitigate erosion during heavy rain storms, was considerably higher at BD sites than at UD sites, and more similar to REF sites.

Without the presence of a well-developed riparian buffer, residential development in shoreland areas often leads to degradation of littoral habitat. For example, a decline in coarse woody structure, which reduces habitat complexity used by many aquatic organisms, a decline of macrophyte populations, which eliminates vital fish and macroinvertebrate habitat, and may interrupt lake ecosystem functioning (Christensen et al. 1996, Jeppesen et al. 1997, Scheffer 1998, Jennings et al. 1999, Elias and Meyer 2003, Jennings et al. 2003, Scheuerell and Schindler 2004, Brauns et al. 2007a). Sediment composition of shallow-water areas may also be affected by shoreland development (Marburg et al. 2006). These impacts may also disrupt habitat coupling interrupting vital ecosystem processes in lakes (Schindler and Scheuerell 2002, Vadeboncoeur et al. 2002).

The expanse and depth of the littoral environment of a lake varies horizontally and vertically with lake geomorphology, distribution of microhabitats, distribution of rooted macrophytes, and water quality, making biotic sampling in this habitat challenging (Wetzel and Likens 2000). Even so, results from our sampling efforts show that the littoral substrate composition (e.g., boulders, cobble, gravel, sand, silt) in the shallow-water benthic habitat varied more across the lakes than within lakes, with East Pond having more boulders at both 0.05 m and 1.0 m transect depths, and East and North Pond having more silt along the 1.0 m transect depth. More silt and sand present in the littoral habitat of North Pond, and more gravel along 0.5 m depth transect of Great Pond, may reflect historic erosion patterns due to degraded riparian buffer areas and the large number of dwellings built near the shorelines of North Pond. When we compared these substrate features across treatments, the only significant difference was that littoral habitats in front of BD and REF properties had less sand along the 0.5 m depth transect than UD properties, perhaps reflecting reduced surface runoff due to the presence of well-developed riparian buffers.

We used an estimate of embeddedness to reflect the extent to which rocks of all sizes were surrounded by sediment or sunken into the lake bottom. An increase in embeddedness can decrease interstitial spaces that provide habitat for juvenile fish and macroinvertebrates, and reduce availability of prey refugia important to sustaining predator-prey interactions (Schlosser 1987, Lynch and Johnson 1989, Beauchamp et al. 1994, Jennings et al. 2003). Percent embeddedness did not differ across the three lakes, suggesting that the causes of embeddedness may operate at similar levels across the three lakes. While differences across treatments were not statistically significant, there was variation with the level of development. Those properties identified as developed without effective buffering capacity (UD) had the highest percent embeddedness at both the 0.5 m and 1.0 m depth transects. High percent embeddedness likely reflects a historic pattern of erosion and sediment loading into the lake in front of poorly buffered or no buffer properties, or properties where residential development is located close to the shoreline. Developed properties with effective buffers (BD) and undeveloped reference sites (REF) had lower levels of embeddedness, supporting the argument that effective riparian buffers function to reduce sediment runoff and help maintain the integrity of the littoral habitat.

There were only a few differences across lakes and treatments for more biological features of the littoral habitats, including the availability of woody structure in all three size categories and aufwuchs cover, features that can serve as indicators of lake ecosystem health. These characteristics were statistically similar for all three lakes and across the three treatments, though REF sites had more woody structure along 0.5 m depth transects compared to BD and UD sites. This may reflect removal patterns by humans at developed sites compared to the absence of human activity at REF sites. Woody structure enhances littoral habitats by contributing to habitat complexity that provides refugia for small fish and macroinvertebrates and serves as a substrate for periphyton (e.g., Vadebencouer and Lodge 2000, Smokorowski and Pratt 2007). Marburg et al. (2006) studied the influence of anthropogenic factors on the presence of littoral coarse wood density in 45 lakes in the Northern Highlands Lake District of Wisconsin, where lake density is high and lakeshore development varies greatly. Overall, these authors found that lakes with highly developed shorelines or lakes with high levels of human activity along parts of the shoreline were lacking coarse woody structure compared to areas with little human disturbance and great variation in amount of woody structure present. Christensen et al. (1996) and Francis and Schindler (2006) report similar findings, but also document a strong positive correlation between riparian forest composition and littoral habitat structure. Aufwuchs cover and density were similar for BD and REF sites at both transect depths, and the values were statistically greater than for those reported for the UD sites along 1.0 m depth transects. These aggregations of microbial lifeforms are a food source for larger organisms, can absorb and remove contaminants in the water column, and serve well as an indicator of water quality (Brauns 2007b). The presence of a well-developed and intact riparian buffer seems to enhance this feature of inshore habitats, helping to mitigate the potential impacts of shoreland development on important littoral habitat features.

The presence of woody structure and aufwuchs supports macroinvertebrate diversity, and the latter two elements of littoral habitat serve as important bioindicators of lake health. In our study, East Pond and North Pond had areas with more medium and coarse woody structure, perhaps helping to explain the higher total number of organisms and mean taxon richness in the rock trap collections than sites on Great Pond. Grab samples taken from littoral habitats indicate East Pond and North Pond, lakes with a higher trophic state index than Great Pond, had roughly two times the total numbers of organisms and greater taxon richness along both transect depths than Great Pond collections. Mean EPT richness in the grab samples increased following a water quality gradient, from poor or below average water quality in East Pond and North Pond to above average water quality in Great Pond, although these differences were not statistically significant. East Pond and North Pond are considered mesotrophic lakes with poor to below average water quality because of shoreline development and external and internal nutrient loading. Such lakes are characterized by high benthic richness and abundance of organisms due, in part, to more complexity and more abundant food resources than less eutrophic lakes.

Differences in grab samples across the three treatment types were limited to the total number of organisms and occurred close to shoreline, along the 0.5 m depth transects where riparian habitat impacts might exert the most influence. BD sites had statistically more total organisms compared to REF and UD sites, suggesting the important influence the presence of a well-developed buffer may have on the littoral macroinvertebrate community of these lakes. Higher aufwuchs cover that provides food and cover for benthic organisms at BD sites may also contribute to this outcome. Our results for macroinvertebrate biodiversity are in line with those from a similar study to compare biotic and abiotic features littoral areas adjacent to reference (undeveloped) and unbuffered developed properties in Vermont lakes (Deeds and Merrell, pers. comm.). There were no statistical differences in species richness or abundance of macroinvertebrates in sandy or rocky littoral communities of reference and unbuffered developed areas. Observers also reported few differences in the relative abundance of macroinvertebrates in the Family Chironomidae, Order Ephemeroptera, or Class Oligochaeta from samples collected from buffered developed and reference sites. Sandy littoral substrate represents a more marginal habitat for macroinvertebrates regardless of lakeshore development status in comparison to rocky littoral areas in this study.

Overall, the results from our surveys of riparian and littoral habitats provide insight into the efficacy of using landscaping best management practices to protect lake ecosystem health. The MSZA standards require a threshold amount of tall vegetation, including trees in each 25 ft^2^ area between the residential structure and the lake, and restrict ground leveling and lawn creation (38 MRSA §435-449). These measures help moderate the erosive power of rain and guard against the formation of impervious surfaces that could facilitate sediment, nutrient, and contaminant runoff that degrades the developed shoreland and adjacent lake ecosystem. Trees along the shoreline not only help protect the shoreline from the erosive power of rain, but also provide shade for both riparian and littoral habitats, and serve as a source of woody structure for near shore littoral habitats. The latter is particularly important to maintain fish and invertebrate populations richness and abundance in the littoral environment.

Our results complement those of Merrell et al. (2013), who found that buffered developed properties along shorelines of five other Maine lakes that met the Maine MSZA standards were not statistically different from undeveloped reference sites for the characteristics that include shading along the shoreline, amount of fine, medium, and course woody structure, amount of deciduous leaf litter, amount of sand, embeddedness, and aufwuchs cover. Not only did our study indicate that well-buffered, developed properties have abiotic and biotic impacts approaching or approximating those of undeveloped properties, but it also suggests that buffered developed properties exert less environmental impact on lake ecosystem health than unbuffered developed properties. Characteristics of unbuffered developed properties were generally worse than reference and buffered developed properties in terms of negative environmental impacts.

Fortunately, lake managers, lake association members, and other stakeholders often focus their efforts to protect lake ecosystem health on these best management practices. The LakeSmart program, one of the largest and most well-known lake protection programs in Maine, uses this approach (Shannon et al. 2017) and participating property owners are quick to recognize both the threat of declining water quality and the potential for successful mitigation strategies (Cole et al. 2017). The current study supports a common premise of lake protection programs like LakeSmart that well-buffered, residential properties can have impacts on the lake ecosystem approaching those of undeveloped properties, and that these properties typically exert less impact on lake ecosystem health than unbuffered residential properties.

Our findings also support expanding the scope of typical lake management efforts to include littoral habitat conservation, perhaps by incorporating the use of macroinvertebrate indicator species to monitor littoral habitat complexity and habitat heterogeneity (Brauns et al. 2007a and b, Poikane et al. 2016, US EPA 2016). Property owners could work to enhance the structural complexity and heterogeneity of disturbed littoral habitat adjacent to their property to protect macroinvertebrate species richness and abundance as well as help assure efficient habitat coupling necessary for a healthy lake ecosystem.

## Acknowledgments

This research was supported in part by the Sustainability Solutions Initiative (National Science Foundation award EPS-0904155 to Maine EPSCoR managed by the University of Maine). We thank faculty colleagues who participated in this five-year, interdisciplinary research project of which our project was a part, including D.W. King (PI), D. Bruesewitz, M. Donihue, J. Fleming, P. Nyhus, T. Reynolds, and S. Dissanayake. Our project also benefitted from technical and logistical support provided by C. Jones, S. Large, E. Arsenault, and A. Pearson. Colby College undergraduate students who provided able and reliable assistance with sample and data collection included T. Abare, E. Arsenault, C. Cummings, M. Davis, S. Doyle, M. Ferguson, A. Junker, S. Judge, S. Large, J. Liang, D. Mealor, N. Moore, S. O’Grady, G-A. Perani, C. Reichler, J. Salay, P. Smithy, W. Supple, M. Susla, B. Timm, S. Weaver, and G. Voigt. D. Homeier, Z. Jaques, S. Madronal, and I. McCullough. A. Schechner provided GIS and data analysis assistance. We are grateful for discussions, advice, and information provided by stakeholders and lake managers including C. Baeder, M. Croft, D. Gay, P. Kallin, M. Shannon, L. Parker, and K. Wall. We gratefully acknowledge the generous support provided by many citizen volunteers affiliated with the Maine Lakes Society, Belgrade Region Conservation Alliance, Belgrade Lakes Association, East Pond Association, North Pond Association, and the staff of the Maine Lakes Resource Center.

---

**Table 2:** Results of ANOVA analysis of sediment organic contents compared across lakes during early season sampling. There was no evidence of interactive effects with time of sampling, and differences were detected only for early season samples.

|  | Sum of Squares | df | Mean Square | F | Significance |
| --- | --- | --- | --- | --- | --- |
| Lake | 4.641 | 2 | 2.320 | 10.622 | 0.001 |
| Residuals | 13.981 | 64 | 0.218 |  |  |

