## Appendix S1 Tables for "The Influence of Shoreline Residential Development on Riparian and Littoral Habitats in the Belgrade Lakes of Maine"

Table S1. A comparison of riparian habitat characteristics among East Pond, North Pond, and Great Pond located in the Belgrade Lakes region of south central Maine, USA. Mean values (± 1 SE) are provided; N = 24 for each characteristic of East Pond and North Pond, and N = 21 for features of Great Pond. An asterisk indicates statistical significance across lakes determined by Kruskal-Wallis test with ties, Holm method adjusted $P < 0.05$. Mean values that do not share the same letter were significantly different at $P < 0.05$ (Wilcoxon rank-sum test).

| Characteristic | East Pond<br>Mean ± SE | North Pond<br>Mean ± SE | Great Pond<br>Mean ± SE |
|---|---|---|---|
| DISTURBANCE INDEX | 1.71 ± 0.17a | 1.92 ± 0.19a | 1.77 ± 0.20a |
| STABILITY INDEX | 2.42 ± 0.15a | 2.54 ± 0.15a | 2.59 ± 0.17a |
| SHADING OVER LAND (%) | | | |
| 1.0 m in from shore | 19.03 ± 1.63a | 17.23 ± 1.73a | 19.47 ± 1.66a |
| SHADING OVER WATER (%) | | | |
| 0.5 m depth | 12.73 ± 1.57a | 11.61 ± 1.59a | 13.79 ± 1.62a |
| 1.0 m depth | 7.18 ± 1.14a | 3.77 ± 0.81a | 8.47 ± 1.55a |
| SLOPE (degrees) | 4.43 ± 0.68a | 9.19 ± 2.22a | 12.23 ± 3.19a |
| BUFFER STRIP (m) | | | |
| Mean width | 6.98 ± 0.82a | 4.49 ± 0.77a | 6.12 ± 0.91a |
| Minimum width | 6.06 ± 0.93a | 4.00 ± 0.83a | 5.19 ± 0.95a |
| Maximum width | 7.54 ± 0.79a | 5.03 ± 0.73a | 7.05 ± 0.96a |
| VEGETATION COVER (%) | | | |
| Tree | 54.58 ± 6.89a | 45.00 ± 7.88a | 61.67 ± 6.89a |
| Tall shrub | 29.00 ± 6.29a | 16.00 ± 4.46a | 29.00 ± 7.24a |
| Low shrub | 36.13 ± 6.51a | 18.42 ± 5.47a | 25.57 ± 6.70a |
| Ground cover* | 73.75 ± 7.42a | 32.67 ± 6.93b | 67.38 ± 8.86a |
| Composite | 7.5 ± 0.504a | 5.21 ± 0.38a | 7.19 ± 0.69a |

| | | | |
|---|---|---|---|
| TREE SIZES (DBH) | | | |
| Regenerating ( < 10 cm) | 24.21 ± 4.42a | 21.13 ± 5.30a | 16.21 ± 2.71a |
| Young (11 - 28 cm) | 6.00 ± 0.98a | 3.83 ± 0.75a | 4.04 ± 0.75a |
| Mature ( > 29 cm) | 2.54 ± 0.56a | 1.79 ± 0.36a | 2.38 ± 0.37a |

Table S2. A comparison of riparian habitat characteristics among the three treatments (Unbuffered developed, Buffered developed, and Reference undeveloped shoreline properties) for East Pond, North Pond, and Great Pond located in the Belgrade Lakes region of south central Maine, USA. An asterisk indicates statistical significance across treatments determined by Kruskal-Wallis test with ties, Holm method adjusted $P < 0.05$. Mean values (± 1 SE) and sample sizes (N) are provided. Mean values that do not share the same letter were significantly different at $P < 0.05$ (Wilcoxon rank-sum test).

| Characteristic | Unbuffered | | Buffered | | Reference | |
|---|---|---|---|---|---|---|
| | N | Mean ± SE | N | Mean ± SE | N | Mean ± SE |
| DISTURBANCE INDEX* | 23 | 2.91 ± 0.06a | 23 | 1.48 ± 0.12b | 24 | 1.04 ± 0.04b |
| STABILITY INDEX* | 23 | 1.91 ± 0.18a | 23 | 2.91 ± 0.06b | 24 | 2.71 ± 0.11b |
| SHADING OVER LAND (%) | | | | | | |
| 1.0 m from shore* | 23 | 11.86 ± 1.82a | 22 | 20.06 ± 1.28b | 24 | 23.55 ± 0.65c |
| SHADING OVER WATER (%) | | | | | | |
| 0.5 m depth* | 23 | 6.93 ± 1.42a | 22 | 12.50 ±1.29b | 24 | 18.31 ± 1.06c |
| 1.0 m depth* | 23 | 3.75 ± 0.94a | 22 | 6.17 ± 1.20ab | 24 | 9.11 ± 1.29b |
| SLOPE (degrees) | 23 | 7.25 ± 1.97a | 22 | 6.95 ± 2.01a | 24 | 10.78 ± 2.58a |
| BUFFER STRIP (m) | | | | | | |
| Mean width* | 23 | 1.31 ± 0.45a | 22 | 7.24 ± 0.71b | 24 | 8.87 ± 0.34b |
| Minimum width* | 23 | 3.65 ± 2.91a | 22 | 6.09 ± 0.87b | 24 | 8.29 ± 0.53c |
| Maximum width* | 23 | 3.12 ± 1.28a | 22 | 8.07 ± 0.63b | 24 | 9.46 ± 0.28b |
| VEGETATION COVER (%) | | | | | | |
| Tree* | 23 | 23.91 ± 5.27a | 22 | 64.54 ± 7.62b | 24 | 71.46 ± 5.73b |
| Tall shrub* | 23 | 5.87 ± 2.30a | 22 | 31.54 ± 6.47b | 24 | 35.83 ± 6.49b |
| Low shrub* | 23 | 11.22 ± 4.09a | 22 | 20.14 ± 5.18a | 24 | 47.71 ± 6.68b |
| Ground cover | 23 | 63.65 ± 8.08a | 22 | 50.45 ± 8.30a | 24 | 58.12 ± 9.15a |
| Composite (unitless)* | 23 | 4.83 ± 0.41a | 22 | 6.95 ± 0.57b | 24 | 8.00 ± 0.50b |
| TREE SIZES, by DBH (%) | | | | | | |
| Regenerating*(< 10 cm) | 24 | 7.83 ± 2.18a | 24 | 21.33 ± 2.87b | 24 | 32.37 ± 5.50b |
| Young* (11 - 28 cm) | 24 | 2.87 ± 0.83a | 24 | 4.08 ± 0.65b | 24 | 6.92 ± 0.85c |
| Mature ( > 29 cm) | 24 | 1.93 ± 0.46a | 24 | 1.87 ± 0.37a | 24 | 2.92 ± 0.46a |

Table S3. A comparison of littoral sediment organics and rock embeddedness recorded for the three study lakes: East Pond, North Pond, and Great Pond located in the Belgrade Lakes region of south central Maine, USA. Mean values (± 1 SE) and sample sizes (N) are provided. An asterisk indicates statistical significance across lakes determined by Kruskal-Wallis test with ties, Holm method adjusted $P < 0.05$. Mean values that do not share the same letter differ at $P < 0.05$ (Wilcoxon rank-sum test).

| Characteristic | East Pond | | North Pond | | Great Pond | |
|---|---|---|---|---|---|---|
| | N | Mean ± SE | N | Mean ± SE | N | Mean ± SE |
| SEDIMENT ORGANICS (%) | | | | | | |
| Early season* | 21 | 3.59 ± 0.70a | 24 | 1.49 ± 0.36b | 24 | 1.09 ± 0.10b |
| Late season | 20 | 3.76 ± 0.80a | 24 | 2.11 ± 0.58a | 22 | 2.08 ± 0.37a |
| EMBEDDEDNESS (%) | | | | | | |
| 0.5 m | 24 | 39.58 ± 6.15a | 24 | 43.05 ± 5.71a | 24 | 32.29 ± 7.24a |
| 1.0 m | 24 | 50.76 ± 7.35a | 24 | 46.96 ± 7.05a | 24 | 53.60 ± 8.13a |

Table S4. A comparison of macroinvertebrate metrics based on grab sample and rock trap collections from the littoral habitat adjacent to study sites on East Pond, North Pond, and Great Pond located in the Belgrade Lakes region of south central Maine, USA. Mean values (± 1 SE) and sample sizes (N) are provided. An asterisk indicates statistical significance across lakes determined by Kruskal-Wallis test with ties, Holm method adjusted $P < 0.05$. Mean values that do not share the same letter differ at $P < 0.05$ (Wilcoxon rank-sum test). Indices are defined in the Methods.

| Characteristic | East Pond | | North Pond | | Great Pond | |
|---|---|---|---|---|---|---|
| | N | Mean ± SE | N | Mean ± SE | N | Mean ± SE |
| Total No. of Organisms | | | | | | |
| 0.5 m | 24 | 35.83 ± 5.38a | 24 | 41.5 ± 7.14a | 23 | 25.52 ± 4.36a |
| 1.0 m | 24 | 53.17 ± 11.11a | 19 | 66.1 ± 10.92a | 22 | 38.54 ± 5.75a |
| Rock trap* | 18 | 97.39 ± 13.48a | 15 | 94.2 ± 10.42a | 14 | 30.29 ± 8.23b |
| Shannon Index | | | | | | |
| 0.5 m | 24 | 4.97 ± 0.37a | 24 | 5.45 ± 0.54a | 23 | 5.27 ± 0.51a |
| 1.0 m | 24 | 4.41 ± 0.33a | 19 | 4.96 ± 0.43a | 24 | 5.58 ± 0.60a |
| Rock trap | 18 | 5.24 ± 0.41a | 15 | 4.77 ± 0.47a | 14 | 5.06 ± 0.53a |
| Mean Richness | | | | | | |
| 0.5 m | 24 | 7.75 ± 0.66a | 24 | 8.79 ± 0.99a | 23 | 7.56 ± 0.80a |
| 1.0 m | 24 | 8.42 ± 0.76a | 19 | 9.79 ± 0.99a | 24 | 8.79 ± 0.82a |
| Rock trap* | 18 | 10.80 ± 0.66a | 15 | 12.0 ± 0.68a | 14 | 7.64 ± 1.07b |
| EPT Richness | | | | | | |
| 0.5 m | 24 | 1.62 ± 0.35a | 24 | 2.08 ± 0.42a | 23 | 2.35 ± 0.36a |
| 1.0 m | 24 | 1.71 ± 0.35a | 19 | 1.95 ± 0.42a | 24 | 2.37 ± 0.36a |
| Rock trap | 18 | 3.00 ± 0.32a | 15 | 3.93 ± 0.36a | 14 | 2.14 ± 0.47a |
| FBI | | | | | | |
| 0.5 m | 24 | 5.81 ± 0.16a | 24 | 5.78 ± 0.20a | 23 | 5.07 ± 0.16a |
| 1.0 m | 24 | 5.51± 0.19a | 19 | 5.75 ± 0.19a | 24 | 5.44 ± 0.16a |

Table S4 (continued)

| Characteristics | East Pond | | North Pond | | Great Pond | |
|---|---|---|---|---|---|---|
| | N | Mean ± SE | N | Mean ± SE | N | Mean ± SE |
| Rock trap* | 18 | 6.54 ± 0.14a | 15 | 4.76 ± 0.14b | 14 | 5.49 ± 0.25c |
| Proportion COTE | | | | | | |
| 0.5 m | 24 | 0.22 ± 0.04a | 24 | 0.21 ± 0.03a | 23 | 0.41 ± 0.06b |
| 1.0 m | 24 | 0.11 ± 0.02a | 19 | 0.12 ± 0.04a | 24 | 0.29 ± 0.05a |
| Rock trap | 18 | 0.31 ± 0.04a | 15 | 0.55 ± 0.06a | 14 | 0.50 ± 0.09a |
| Proportion Crustacea and Mollusca | | | | | | |
| 0.5 m | 24 | 0.31 ± 0.05a | 24 | 0.27 ± 0.04a | 23 | 0.16 ± 0.04a |
| 1.0 m | 24 | 0.44 ± 0.06a | 19 | 0.49 ± 0.05a | 24 | 0.37 ± 0.05a |
| Rock trap | 18 | 0.12 ± 0.04a | 15 | 0.28 ± 0.06a | 14 | 0.16 ± 0.05a |
| Proportion Chironomidae | | | | | | |
| 0.5 m | 24 | 0.21 ± 0.05a | 24 | 0.25 ± 0.05a | 23 | 0.17 ± 0.03a |
| 1.0 m | 24 | 0.12 ± 0.02a | 19 | 0.14 ± 0.03a | 24 | 0.10 ± 0.02a |
| Rock trap* | 18 | 0.37 ± 0.05a | 15 | 0.06 ± 0.02b | 14 | 0.02 ± 0.01c |

Table S5. A comparison of littoral sediment organics and rock embeddedness across the three treatment types: Unbuffered developed, Buffered developed, and Reference undeveloped shoreline properties for sites on East Pond, North Pond, and Great Pond located in the Belgrade Lakes region of south central Maine, USA. Mean values (± 1 SE) and sample sizes (N) are provided. An asterisk indicates statistical significance across treatments determined by Kruskal-Wallis test with ties, Holm method adjusted *P* < 0.05. Mean values that do not share the same letter differ at *P* < 0.05 (Wilcoxon rank-sum test).

| Characteristic | Unbuffered | | Buffered | | Reference | |
|---|---|---|---|---|---|---|
| | N | Mean ± SE | N | Mean ± SE | N | Mean ± SE |
| SEDIMENT ORGANICS (%) | | | | | | |
| Early season | 24 | 1.88 ± 0.40a | 21 | 1.84 ± 0.42a | 22 | 2.34 ± 0.65a |
| Late season | 24 | 2.33 ± 0.40a | 22 | 1.79 ± 0.38a | 20 | 3.81 ± 0.94a |
| EMBEDDEDNESS (%) | | | | | | |
| 0.5 m | 24 | 55.13 ± 6.96a | 24 | 25.66 ± 4.60a | 24 | 34.13 ± 5.97a |
| 1.0 m | 24 | 66.29 ± 7.37a | 24 | 41.90 ± 6.68a | 24 | 43.18 ± 7.44a |

Table S6. A comparison of littoral structural characteristics among Unbuffered developed, Buffered developed, and Reference undeveloped shoreline properties on East Pond, North Pond, and Great Pond located in the Belgrade Lakes region of south central Maine, USA. Mean values (± 1 SE) and sample sizes (N) are provided. An asterisk indicates statistical significance across treatments determined by Kruskal - Wallis test with ties, Holm method adjusted $P < 0.05$. Mean values that do not share the same letter differ at $P < 0.05$ (Wilcoxon rank-sum test).

| Characteristics | Unbuffered | | Buffered | | Reference | |
|---|---|---|---|---|---|---|
| | N | Mean ± SE | N | Mean ± SE | N | Mean ± SE |
| FINE WOODY STRUCTURE (mean #/quadrat) | | | | | | |
| 0.5 m | 24 | 1.18 ± 0.30a | 24 | 2.59 ± 0.55a | 24 | 3.38 ± 0.73a |
| 1.0 m | 24 | 1.70 ± 0.62a | 24 | 1.40 ± 0.67a | 24 | 1.72 ± 0.45a |
| MEDIUM WOODY STRUCTURE (mean #/quadrat) | | | | | | |
| 0.5 m | 24 | 0.20 ± 0.05a | 24 | 0.46 ± 0.17a | 24 | 1.06 ± 0.28a |
| 1.0 m | 24 | 0.29 ± 0.09a | 24 | 0.19 ± 0.06a | 24 | 0.89 ± 0.51a |
| COARSE WOODY STRUCTURE (mean #/quadrat) | | | | | | |
| 0.5 m* | 24 | 0.05 ± 0.02a | 24 | 0.11 ± 0.03a | 24 | 0.37 ± 0.10b |
| 1.0 m | 24 | 0.11 ± 0.04a | 24 | 0.09 ± 0.03a | 24 | 0.15 ± 0.04a |
| LEAF LITTER (% transect) | | | | | | |
| 0.5 m | 24 | 5.07 ± 1.93a | 24 | 2.49 ± 0.70a | 24 | 12.17 ± 5.40a |
| 1.0 m | 24 | 9.29 ± 3.34a | 24 | 3.26 ± 1.30a | 24 | 1.04 ± 0.46a |
| AUFWUCHS COVER (rating 1-10) | | | | | | |
| 0.5 m | 24 | 3.78 ± 0.67a | 24 | 5.42 ± 0.59a | 24 | 5.77 ± 0.69a |
| 1.0 m* | 24 | 3.32 ± 0.67a | 24 | 6.83 ± 0.54b | 24 | 6.89 ± 0.61b |

Table S6 (continued)

| Characteristic | Unbuffered | | Buffered | | Reference | |
|---|---|---|---|---|---|---|
| | N | Mean ± SE | N | Mean ± SE | N | Mean ± SE |
| AUFWUCHS DENSITY (rating 1-3) | | | | | | |
| 0.5 m | 24 | 0.97 ± 0.18a | 24 | 1.27 ± 0.13a | 24 | 1.42 ± 0.17a |
| 1.0 m | 24 | 0.95 ± 0.19a | 24 | 1.46 ± 0.16a | 24 | 1.59 ± 0.20a |
| MACROPHYTE ABUNDANCE (% transect) | | | | | | |
| 0.5 m | 23 | 9.61 ± 3.24a | 23 | 6.91 ± 3.35a | 24 | 3.90 ± 1.62a |
| 1.0 m | 22 | 14.11 ± 4.96a | 23 | 10.73 ± 4.56a | 24 | 5.22 ± 1.71a |

Table S7. A comparison of macroinvertebrate metrics for grab sample and rock trap collections from the littoral habitat adjacent to Unbuffered developed, Buffered developed, and Reference undeveloped properties on East Pond, North Pond, and Great Pond located in the Belgrade Lakes region of south central Maine, USA. Mean values (± 1 SE) and sample sizes (N) are provided. An asterisk indicates statistical significance across treatments determined by Kruskal - Wallis test with ties, Holm method adjusted $P < 0.05$. Mean values that do not share the same letter differ at $P < 0.05$ (Wilcoxon rank-sum test). Indices are defined in the Methods.

| Characteristic | Unbuffered | | Buffered | | Reference | |
|---|---|---|---|---|---|---|
| | N | Mean ± SE | N | Mean ± SE | N | Mean ± SE |
| Total No. of Organisms | | | | | | |
| 0.5 m* | 24 | 28.37 ± 7.33a | 24 | 48.21 ± 4.85b | 23 | 26.30 ± 3.79a |
| 1.0 m | 23 | 64.91 ± 11.97a | 22 | 50.36 ± 7.17a | 22 | 38.91 ± 8.26a |
| Rock trap | 18 | 70.72 ± 11.13a | 17 | 73.47 ± 11.3a | 12 | 89.00 ± 20.94a |
| Shannon Index | | | | | | |
| 0.5 m | 24 | 4.70 ± 0.47a | 24 | 5.92 ± 0.45a | 23 | 5.07 ± 0.49a |
| 1.0 m | 23 | 5.39 ± 0.47a | 22 | 5.27 ± 0.54a | 22 | 4.28 ± 0.40a |
| Rock trap | 18 | 5.55 ± 0.48a | 17 | 4.75 ± 0.34a | 12 | 4.67 ± 0.57a |
| Mean Richness | | | | | | |
| 0.5 m* | 24 | 6.50 ± 0.78a | 24 | 10.25 ± 0.80b | 23 | 7.35 ± 0.71a |
| 1.0 m | 23 | 9.87 ± 0.86a | 22 | 9.64 ± 0.79a | 22 | 7.27 ± 0.80a |
| Rock trap | 18 | 11.33 ± 0.89a | 17 | 9.29 ± 0.63a | 12 | 9.92 ± 1.23a |
| EPT Richness | | | | | | |
| 0.5 m | 24 | 1.46 ± 0.38a | 24 | 2.71 ± 0.42a | 23 | 1.87 ± 0.28c |
| 1.0 m | 23 | 2.26 ± 0.40a | 22 | 2.14 ± 0.41a | 22 | 1.64 ± 0.30a |
| Rock trap | 18 | 3.44 ± 0.41a | 17 | 2.88 ± 0.37a | 12 | 2.67 ± 0.45a |
| FBI | | | | | | |
| 0.5 m | 24 | 5.39 ± 0.22a | 24 | 5.66 ± 0.16a | 23 | 5.64 ± 0.18a |

Table S7 (cont.)

| Characteristic | Unbuffered | | Buffered | | Reference | |
|---|---|---|---|---|---|---|
| | N | Mean ± SE | N | Mean ± SE | N | Mean ± SE |
| 1.0 m | 23 | 5.74 ± 0.16a | 22 | 5.48 ± 0.19a | 22 | 5.43 ± 0.19a |
| Rock trap | 18 | 5.63 ± 0.22a | 17 | 5.35 ± 0.24a | 12 | 6.14 ± 0.31a |
| % COTE | | | | | | |
| 0.5 m | 24 | 0.27 ± 0.06a | 24 | 0.26 ± 0.04a | 23 | 0.29 ± 0.04a |
| 1.0 m | 23 | 0.14 ± 0.03a | 22 | 0.16 ± 0.03a | 22 | 0.24 ± 0.06a |
| Rock trap | 18 | 0.41 ± 0.05a | 17 | 0.53 ± 0.06a | 12 | 0.36 ± 0.08a |
| % Crustacea and Mollusca | | | | | | |
| 0.5 m | 24 | 0.22 ± 0.04a | 24 | 0.25 ± 0.05a | 23 | 0.28 ± 0.05a |
| 1.0 m | 23 | 0.40 ± 0.05a | 22 | 0.45 ± 0.06a | 22 | 0.43 ± 0.06a |
| Rock trap | 18 | 0.21 ± 0.05a | 17 | 0.18 ± 0.04a | 12 | 0.17 ± 0.06a |
| % Chironomidae | | | | | | |
| 0.5 m | 24 | 0.15 ± 0.05a | 24 | 0.27 ± 0.04a | 23 | 0.21 ± 0.04a |
| 1.0 m | 23 | 0.10 ± 0.02a | 22 | 0.13 ± 0.03a | 22 | 0.13 ± 0.02a |
| Rock trap | 18 | 0.15 ± 0.05a | 17 | 0.14 ± 0.05a | 12 | 0.22 ± 0.08a |
